# Pan-cancer identification of clinically relevant genomic subtypes using outcome-weighted integrative clustering

**DOI:** 10.1101/2020.05.11.084798

**Authors:** Arshi Arora, Adam B. Olshen, Venkatraman E. Seshan, Ronglai Shen

**Affiliations:** Department of Epidemiology and Biostatistics, Memorial Sloan Kettering Cancer Center, New York, NY; Department of Epidemiology and Biostatistics, University of California at San Francisco, CA; Helen Diller Family Comprehensive Cancer Center, University of California at San Francisco, CA

**Author notes:** Corresponding authors Correspondence to Ronglai Shen.

**Keywords:** Integrative Genomics, Supervised Clustering, Cancer Genomics, Statistical Methods, Data Integration

## Abstract

Molecular phenotypes of cancer are complex and influenced by a multitude of factors. Conventional unsupervised clustering of heterogeneous cancer patient populations is inevitably driven by the dominant variation from major factors such as cell-of-origin or histology. Drawing from ideas in supervised text classification, we developed survClust, an outcome-weighted clustering algorithm for integrative patient stratification. We show survClust outperforms unsupervised clustering in identifying cancer patient subpopulations characterized by specific genomic phenotypes with more aggressive clinical behavior. The algorithm and tools we developed have direct utility toward clinically relevant patient stratification based on tumor genomics to inform clinical decision-making.

## INTRODUCTION

Cancer is a complex disease with heterogeneous clinical outcomes. Understanding how patients respond to treatment and what drives disease progression and metastasis is critical for managing and curing the disease. Linking comprehensive molecular profiling data with patient outcome carries great promise in addressing such important clinical questions. This requires innovative statistical and computational methods designed for integrative analysis of multidimensional data sets to model intra-tumor and inter-patient heterogeneity at genomic, epigenetic, and transcriptomic levels. Each of these molecular dimensions is correlated yet characterize the disease in their own unique way. In order to arrive at a comprehensive molecular portrait of the tumor, multiple groups have proposed statistical and computational algorithms to synthesize various channels of information including methods developed by us (iCluster^1,2^) and others (PARADIGM^3^, CoCA^4^, SNF^5^, CIMLR^6^) to stratify disease populations. However, the majority of the work has focused on unsupervised clustering, utilizing the molecular data alone.

Unsupervised learning does not necessarily lead to unique answers as the data are often complex and multi-faceted. Consider the problem of clustering a collection of documents in text mining where multiple structures can be present including authorship, topic, and style. The outcome of the clustering is likely driven by a mixture of these underlying structures. As a result, there is often no single “right” answer in unsupervised clustering problems. In most complex data applications, many local optima exist that poses special challenges in optimization. Xing et al.^7^ proposed a weighted distance metric allowing users to specify what they consider “meaningful” in defining similarity toward a more efficient and local-optima free clustering performance.

Drawing analogy with the text learning problem described above, the molecular profile of a tumor is influenced by a multitude of factors including tissue-of-origin^8^, histology (e.g., squamous vs. adenocarcinoma), tumor microenvironment (e.g., immune cell infiltration^9^), dedifferentiation states^10^, and specific pathway activation^11^. Conventional unsupervised clustering applied to the most variable features is inevitably driven by the dominant variation from major factors, for example, cell-of-origin^8^ or ancestry^12^ (germline variation) in the study cohort. When patient outcome related stratification is of interest, a more directed clustering approach is needed.

We present *survClust*, an outcome-weighted integrative clustering algorithm for survival stratification based on multi-dimensional omics-profiling data. The algorithm learns a weighted distance matrix that down-weights molecular features with no relevance to the outcome of interest. This method can be used on individual platforms alone, or by integrating various molecular platforms, to mine biological information leading to distinct survival subgroups. We analyzed over 6,000 tumors across 18 cancer types. Each disease type was classified by *survClust*, based on six molecular assays – somatic point mutations, DNA copy number, DNA methylation, mRNA expression, miRNA expression, protein expression, and the integration of the six assays. The results have revealed novel survival subtypes not previously identified by unsupervised clustering.

## RESULTS

### The *survClust* model: motivation and method overview

The molecular profile of a tumor often harbors information on a multitude of factors including cell lineage, tumor microenvironment, cell differentiation and other clinical and histopathological features. Some of these factors are associated with treatment response and/or survival outcome, while others are not. If a particular patient outcome (e.g., patient survival) is of interest, a more supervised approach is needed. We demonstrate this using a simulated data example (**Fig. 1a, Supplementary Fig 1**). In this scenario, we simulated three risk subgroups in a cohort of 300 hypothetical patient samples with distinct survival hazard rates in each subgroup (a median survival of 4, 3, and 2 years respectively). A set of 15 features was then simulated from a mixture Gaussian distribution with different means in the three risk subgroups. Another set of 15 features was simulated in the same way but permutated to disrupt the feature-risk group association. A third group of 270 features were simulated from Gaussian noise. Figure 1b shows that an unsupervised clustering using the K-means algorithm failed to identify the survival subtypes in the context of complex feature variations. To identify outcome-associated clustering solution, *survClust* utilizes a weighted distance metric:

**Figure 1:**
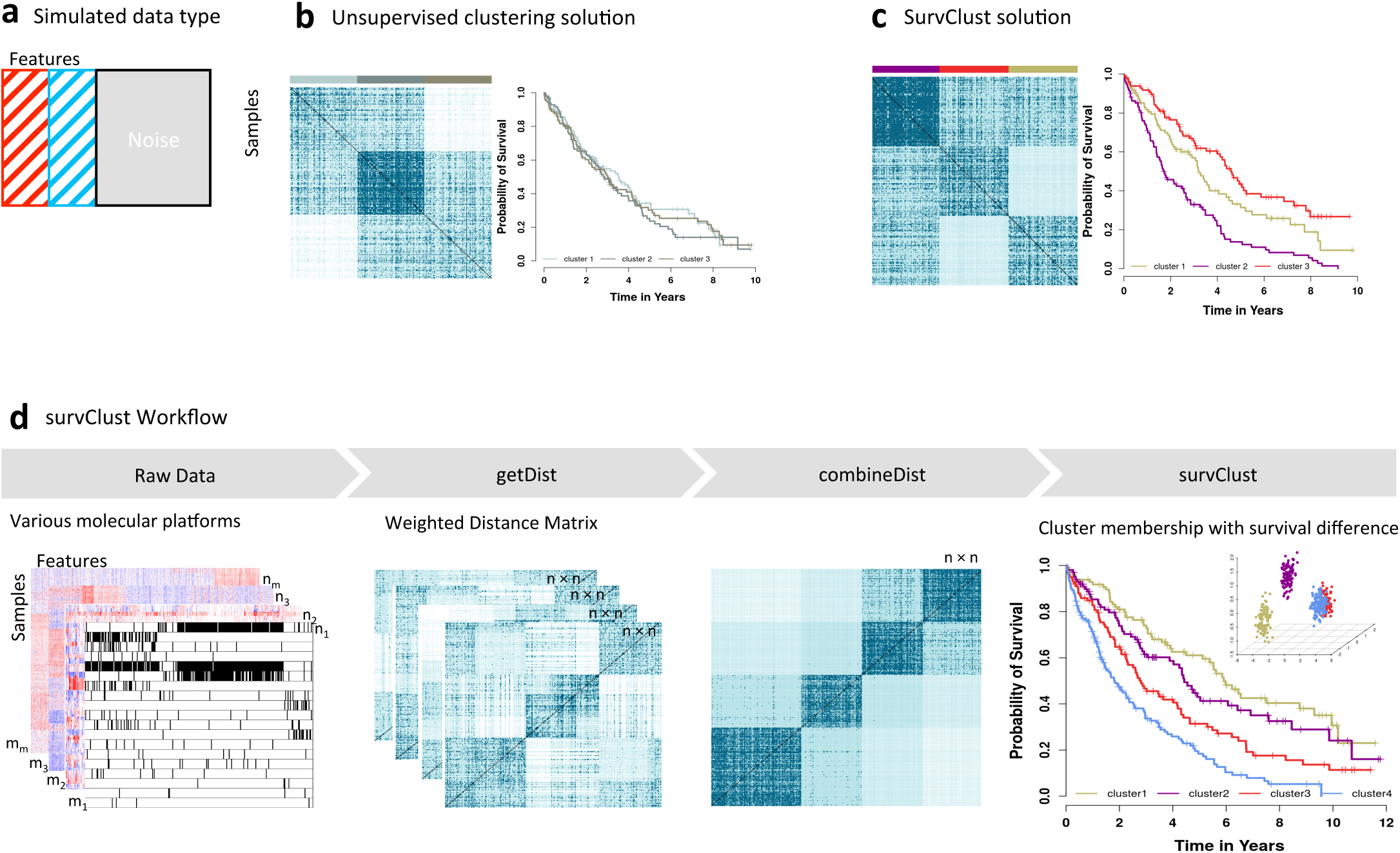
Overview of *survClust.* (a) A simulated data example, consisting of features that define 3 patient subtypes without direct association with survival (shaded in red), features that define 3 patient subtypes with distinct survival outcome (shaded in blue), and random features generated from Gaussian noise (grey). *(b)* Euclidean distance matrix demonstrating patient-level pairwise similarity, with darker blue shade representative of higher similarity. Color panels above the distance matrix show the three class solution obtained by unsupervised algorithm via k-means and the concordance between the simulated 3 survival subtypes (the truth). Kaplan Meier curves for the 3 unsupervised subtypes show no distinction in survival outcome. *(c) survClust* employs a patient outcome weighted distance matrix to identify the desired subtypes with distinct Kaplan Meier curves. *(d) survClust* allows integrative analysis of multiple data modalities to identify survival-associated molecular subtypes.

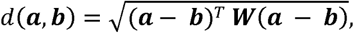

where (***a***,***b***) denote a pair of sample vectors measured for *p* features, and ***W*** is a diagonal weight matrix over *p* features with ***w*** = *diag* {*w*_l_, …,*w*_*p*_}. The weights *w*_p_ ′*s*, are obtained by fitting a univariate cox proportional hazards model for each feature in the training data with repeated training-test sample splits for cross-validation (see more details in the Methods Section). Figure 1c shows that *survClust* was able to identify the true risk groups with 97.15% accuracy [95% CI = 94% - 100%], whereas the accuracy from an unsupervised clustering was 67.50% without reducing the effect of the survival unrelated and noise features.

Our algorithm allows the integration of multiple data modalities. Given *m* data types measured over respective feature space (**Fig. 1d**), the algorithm learns a weighted distance matrix from each molecular data type incorporating a vector of Cox regression hazard ratio as weights. Each feature is weighed and a pairwise distance matrix is calculated (we refer to this step as **getDist)**. This step reduces the computation space considerably by transforming the problem from sample by feature to sample by sample. Note that, different sample sizes across data types are allowed, i.e., a sample can be measured for some but not all platforms. Next, the weighted pairwise distance matrices are integrated by summing over weighted *m* data types (**combineDist**), which retains all samples with at least one data type available, with complete pairwise information. **survClust** then projects the integrated and weighted distance matrix into a lower dimensional space via multidimensional scaling (MDS) and then clusters sample points into subgroups via the K-means algorithm. More details can be found in the Methods Section.

### *survClust* is more powerful than unsupervised clustering in identifying clinically relevant molecular subtypes

We applied *survClust* to the TCGA data set including 6,209 tumor samples in 18 cancer types to identify survival outcome-associated subtypes defined by somatic mutation, DNA copy number, DNA methylation, mRNA expression, and protein expression, individually and integratively. A summary of the sample sizes and feature space is included in Supplementary Table 1. Supplementary Table 2 compares the survival association (log-rank statistic) for the *survClust* integrated subtypes versus those derived from unsupervised clustering methods commonly used in TCGA studies including COCA and iCluster. The log-rank statistic compares estimates of the hazard functions of each subgroup comparing to the expected values under the null hypothesis (all subgroups have identical hazard functions). Larger log-rank statistic suggests stronger evidence of survival association. By differentially weighting the molecular features by the corresponding survival association in constructing the distance matrix, we show that *survClust* is more powerful in identifying subtypes that are directly relevant to stratify the outcome of interest, leading to substantially more distinct survival subgroups than those existing molecular subclasses obtained by unsupervised clustering. To further demonstrate, we highlight the *survClust* analysis of low-grade glioma and kidney papillary renal cell carcinoma below.

### *survClust* identifies a poor prognostic *IDH-*mutant low-grade glioma subgroup

Low Grade Gliomas (LGG) have a unique molecular footprint, characterized by *IDH1/2* mutation status and co-deletion in chromosome 1p and19q regions of the genome^13^. As shown previously, mutations in *IDH1* and *IDH2* genes are present in a majority of the low-grade gliomas and define a subtype associated with favorable prognosis^14^. *IDH*-mutant tumors with chromosome 1p and 19q codeletion (IDHmut-codel) exhibit the most prolonged survival times followed by *IDH*-mutant tumors without the codeletion (IDHmut-non-codel), with *IDH-*wt tumors demonstrating more aggressive clinical behavior. We performed *survClust* on 6 available molecular platforms (somatic mutation, DNA copy number, DNA methylation, mRNA expression, and protein expression) in 512 LGG samples as profiled by the TCGA. The optimal number of clusters *k* was chosen by assessing *survClust* fits over log-rank test statistics and standardized pooled within-cluster sum-of-squares in cross-validation (see Methods Section). Cross-validation was performed to ensure unbiased estimation of survival association and to avoid over-fitting.

The integrated *survClust* solution for LGG was optimized at k=5, with the *IDH*-mutant-codel (c3) and *IDH*-mutant-non-codel (c1) subtypes associated with good prognosis as expected (**Fig 2a**). By contrast, the *IDH*-wt subclass (c5) showed association with poor survival, enriched for mutations in *EGFR* and *PTEN* gene and concurrent chromosome 7 gain and 10 loss, resembling glioblastomas. Interestingly, *survClust* identified a small *IDH*-mutant subtype characterized by *CDKN2A* deletion (c4), which showed markedly worse survival among the *IDH*-mutant tumors, similar to the *IDH*-wt group (c5) that tends to behave far more aggressively with prognosis similar to glioblastomas. In addition, a copy number quiet subgroup (c2) was identified, showing high expression of mir-1307 and mir-29c (**Supplementary Fig 3**). These results highlight the strength of *survClust* in identifying clinically relevant molecular stratifications and the potential to refine the existing paradigm in glioma subtyping to inform clinical decision-making.

**Figure 2:**
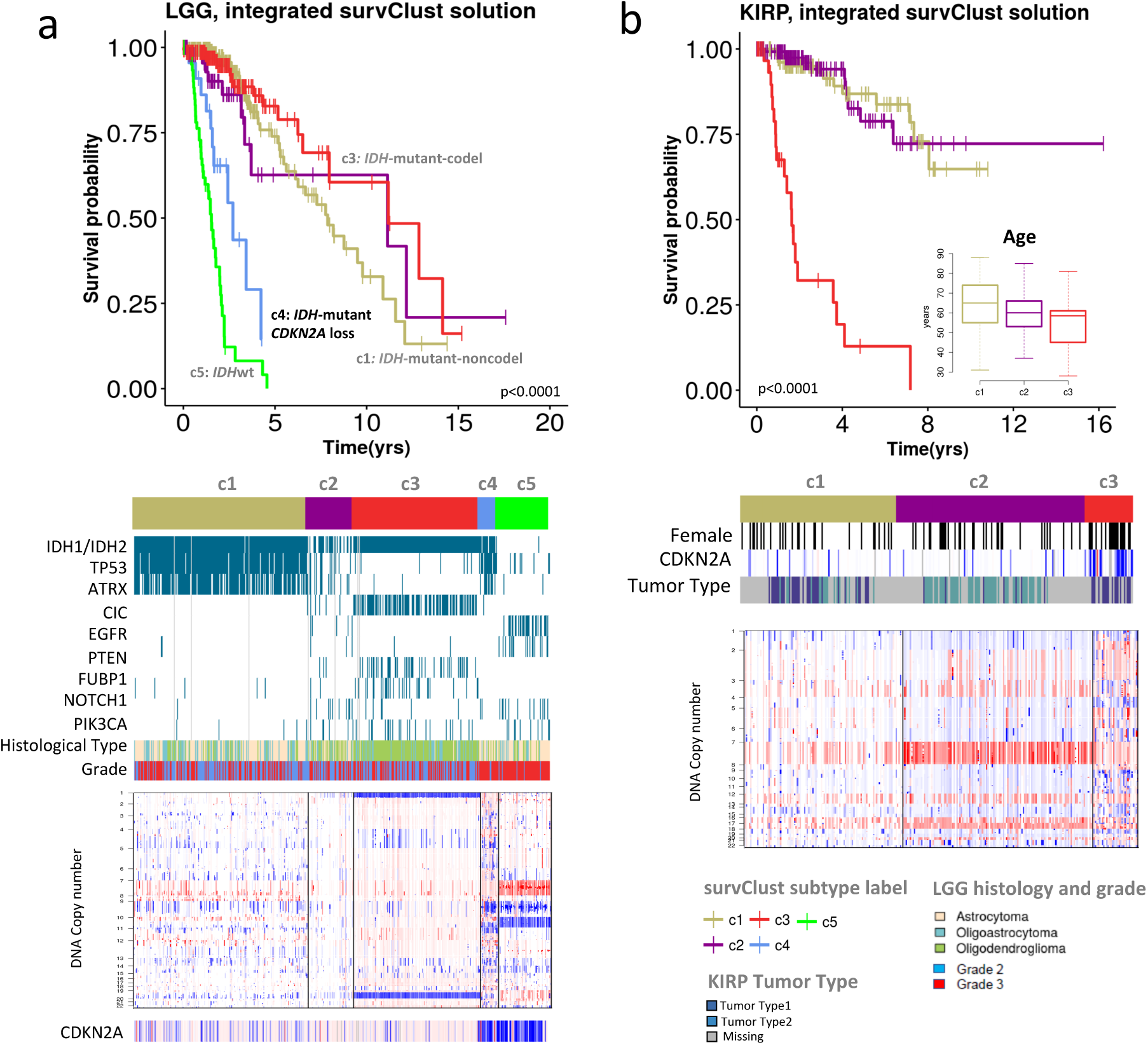
Outcome-weighted integrative clustering of low grade glioma and kidney papillary cell carcinoma using *survClust*. *(a) survClust* identifies an *IDH-*mutant *CDKN2A*-loss subtype similar to *IDH*-wt tumors in terms of aggressive clinical behavior. Top: Kaplan-Meier curves of the integrated *survClust* subtypes of LGG. Middle: *Panelmap* summarizing major association of mutational and clinical features of the integrated LGG subtypes. Bottom: Copy number profile for each of the integrated subtypes. *(b) survClust* identifies prognostic kidney papillary renal cell carcinoma (KIRP) subtypes.

### *survClust* identifies prognostic subtypes of kidney papillary renal-cell carcinoma (KIRP)

Three survival distinct subtypes were identified using *survClust* integrating DNA copy number, mRNA expression, DNA methylation, miRNA and protein expression assay profiled in 289 tumor samples. The c3 subtype was associated with poor survival (median survival time = 1.63 yrs) (**Fig 2b**), associated with younger age (median age 57 yrs) and more female gender (55%). The defining genomic characteristics include *CDKN2A* loss, arm-level gains in multiple chromosomes including 7, 12, 15 and 17 as described previously^15^.

### *survClust* identifies clinically relevant mutational subgroups across cancer types

*survClust* is a flexible framework and can be applied to individual data types for patient stratification. For example, somatic mutation based stratification is often of interest in a clinical sequencing setting. To illustrate that, we applied *survClust* to mutation data alone using a hazard ratio weighted binary distance-based clustering. A *circomap* plot was created to facilitate annotation and visualization of the results across cancer types (**Fig 3a**). *survClust* identified high TMB subgroups in nearly all cancer types included in this analysis. Correlating mutational signatures^16^ with these subtypes in the *circomap* plot further revealed etiology underlying these hypermutated tumors. The smoking signature tracks lung cancer (LUSC and LUAD) and the subset of head and neck cancer (HNSC) with elevated TMB. The DNA mismatch repair (MMR) signature tracks high TMB subgroups in stomach (STAD), endometrial cancer (UCEC), and colon cancer (COAD). The APOBEC signature is prevalent in bladder (BLCA) and cervical cancers (CESC). Finally, the aristolochic acid signature (signature 22) is enriched in a liver cancer subgroup identified by *survClust* (**Supplementary Fig 4e**), which is consistent with aristolochic acid and their derivatives being implicated in liver cancers in Asian populations^17^.

**Figure 3:**
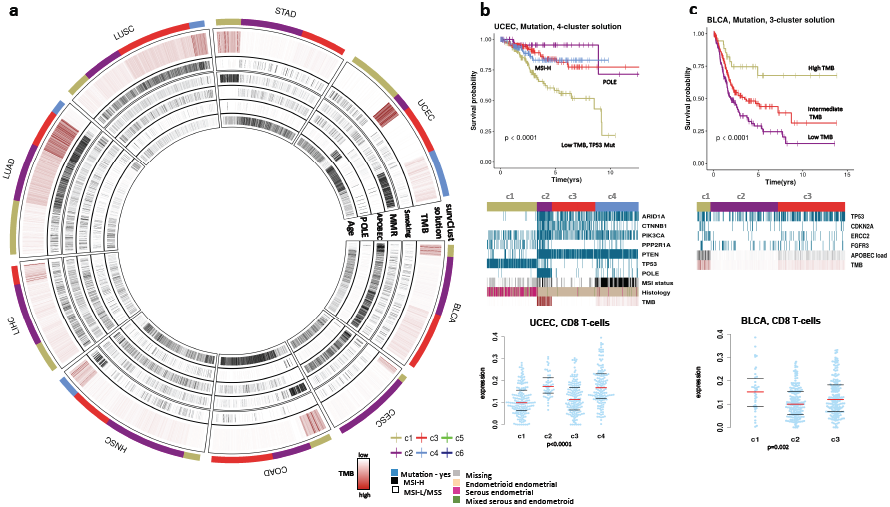
*survClust* identifies mutational subtypes associated with survival across cancer types. *(a) Circomap* showing total mutation burden (TMB) in brown color and mutational signatures (smoking, MMR, APOBEC, POLE and aging) in tumors across bladder (BLCA), cervical (CESC), colon (COAD), head and neck (HNSC), liver (LIHC), lung adenocarcinoma (LUAD), lung Squamous Cell (LUSC), stomach (STAD), and endometrial (UCEC) cancers. Outer circle indicates mutation-based *survClust* membership. *(b) survClust* mutation subtypes in endometrial cancer. From top to bottom: Kaplan-Meier curves for the 4 mutation subtypes, *panelmap* depicting significantly mutated genes, MSI status, Histology and TMB associated with the subtypes, and beeswarm plot showing CD8 T-cell marker expression (y-axis) across the 4 subtype (x-axis). Red line depicts the median, and top and bottom black bars represent the 25^th^ and 75^th^ percentile respectively. *(c) survClust* mutation subtypes in bladder cancer. From top to bottom: Kaplan-Meier curves for the 3 mutation subtypes, *panelmap* depicting significantly mutated genes, Papillary histology (yes – black, no-white), APOBEC load and TMB associated with the 3 subtypes, and beeswarm plot showing CD8 T-cell expression (y-axis) across the 3 subtypes (x-axis).

In endometrial cancer, *survClust* confirmed a previously known ultra-high mutated subtype associated with the POLE mutation signature (c2) and a hypermutated microsatellite instability (MSI) (c4) subtype^18^ (**Fig 3b**). The *panelmap* in Figure 3b (middle panel) shows that c4 correlated well with clinical MSI status (P<0.001) and predominantly carried mutants in *ARID1A, PIK3CA* and *PTEN* genes. The c1 subtype, consisting of primarily high-grade serious tumors, was associated with worse outcome with a 5-year survival of 58% compared to 95%, 84%, and 83% for c2 (POLE), c3, and c4 (MMR) respectively, and characterized by higher frequency of mutations in *TP53, PPP2R1A* genes, low TMB and older patients with serous endometrial tumors (60%). The c3 subtype was characterized by higher frequency of *CTNNB1* mutants. Immune cell decomposition data derived using the CIBERSORT^19^ algorithm was also correlated with the subgroups. Interestingly, high expression of CD8 T-cell immune marker was observed in the POLE (c2) and MSI (c4) subtype (P < 0.001) (**Fig 3b**).

*survClust* stratified the bladder cancer cohort into 3 TMB subgroups – with high (c1), intermediate (c3) and low (c2) mutation burden. The c1 subtype was associated with good outcome, high TMB, high neoantigen load, high APOBEC load, and high expression of the CD8 T-Cell immune marker (P=0.002) (**Fig 3c**). The c3 subtype showed intermediate TMB and APOBEC load with a median survival time of 3.48 yrs. Patients with a low TMB and low APOBEC load performed the worst in terms of survival with a median survival time of 1.91 yrs.

A similar pattern emerged when *survClust* was run on colorectal cancer mutation data classifying the disease population into three clusters – two low TMB groups and a MMR-associated high TMB group (c1) (**Supplementary Fig 4b**). c1 was also associated with CD8 T-cell infiltration (P = 0.004) and showed concordance with MLH1 silencing status. A similar subdivision of low TMB group by *TP53* mutation status was seen where c3 carried *TP53* mutant samples unlike c2. Correlation with histology revealed significant enrichment of mucinous adenocarcinoma subtype in c1 and c2 (c1, n=20, 29%; c2, n=24, 20%) compared to c3 (n=9, 5%). In addition to the hypermutated subtypes of endometrial, bladder and colorectal cancers, we also observed high TMB subgroups with concurrently high expression of CD8 T-cell markers in cervical cancer c1 subtype (**Supplementary Fig 4a and 5a**), head and neck cancer c4 subtype **(Supplementary Fig 4c, 5c)**, lung adenocarcinoma c3 subtype (**Supplementary Fig 4f and 5f**), lung squamous cell carcinoma c4 subtype (**Supplementary Fig 4g and 5g**) and stomach cancer c1 subtype (**Supplementary Fig 4h and 5h**). There are prior observations that high mutational burden is associated with increased neo-antigen load and activated T-cell infiltration in lung cancer^20^. Our analysis revealed that such association may be more widely present in multiple cancer types.

### *survClust* identifies distinct copy number subtypes associated with clinical features across cancer types

To identify copy number alterations that define clinically relevant subtypes, segmented data of 18 cancer types was processed via the CBS algorithm^21^ and analyzed with *survClust*. Subtypes characterized by different degrees in the Fraction of Genome Altered (FGA) emerged in various cancer types **(Fig 4)**. Interestingly, low FGA was associated with better survival in several cancer types including colon, head and neck, lung adenocarcinoma, soft tissue sarcoma and endometrial cancer (**Supplementary Fig 6 and 7).**

**Figure 4:**
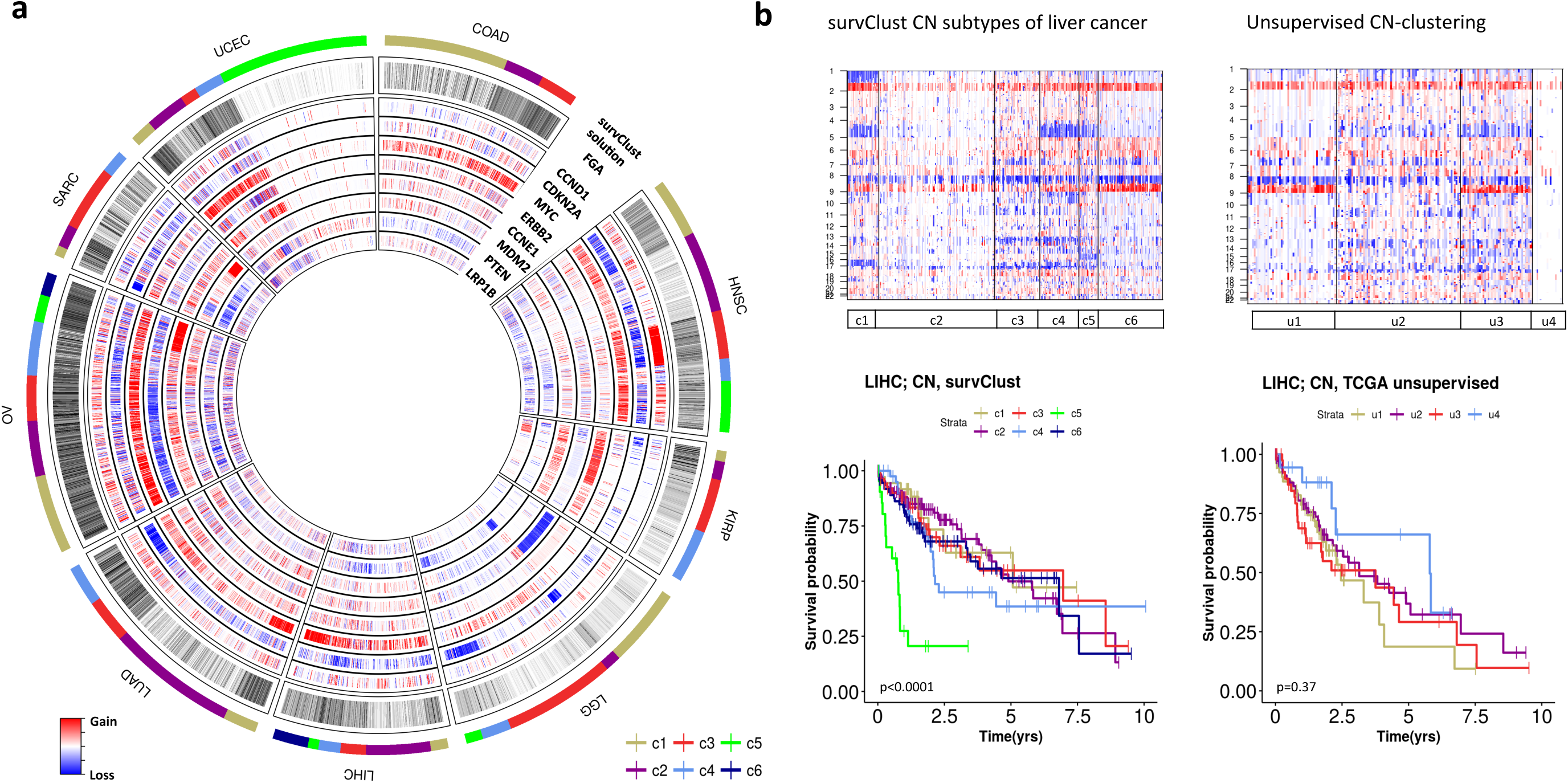
*survClust* identifies copy number paQerns associated with patient survival outcome across various cancer types. *(a) Circomap* showing fraction genome altered (FGA) and gene level copy number alterations in each tumor across colorectal (COAD), head and neck (HNSC,) kidney renal papillary cell carcinoma (KIRP), low grade glioma (LGG), liver (LIHC), lung adenocarcinoma (LUAD), ovarian (OV), sol-tissue sarcoma (SARC) and endometiral (UCEC) cancers. Outer circle indicates the *survClust* membership. *(b) survClust* is more powerful than unsupervised clustering in identifying survival-associated copy number subtypes in liver cancer.

The *circomap* plot in Figure 4a also revealed association of subtypes with high-level amplification of major cancer genes including *CCND1* amplification in head and neck cancer (c3), *CCNE1* (c5) and *AKT2*(c6) amplification in ovarian cancer, and *MDM2* amplification (c4) in sarcoma (**Supplementary Fig 6**). Notably, amplification of 19q13.2 region in ovarian cancer c6 subtype harboring the *AKT2* gene is associated with poor survival (**Supplementary Fig 7f, Supplementary Table 8**) which was consistent with previous findings that *AKT2* amplification is associated with ovarian cancer aggressiveness^22^. *CCND1* amplified subtype of head and neck cancer (c3) was also associated with poor survival (**Supplementary Fig 7b**). Amplification in the *MYC* gene is broadly present in multiple cancer types (**Fig 3a** *circomap*). Among cancer gene deletions, *CDKN2A* loss was observed to define multiple subgroups associated with poor survival including papillary kidney cancer (c1), low-grade glioma (c4), lung adenocarcinoma (c4), and soft tissue sarcoma (c1) (**Supplementary Fig 6 and 7**).

Colorectal cancer was classified into three varying FGA subtypes with prognostic implications. c1 had low FGA and, c2 and c3 carried heavy genome alterations (**Supplementary Fig 6a**). Even though c1 and c2 had dissimilar FGA, they performed similar in terms of survival as compared to c3, which had poor outcome with median survival time of 4.5 yrs. (**Supplementary Fig 7a**). Gain in the MYC gene was seen throughout the cancer type and c2 was uniquely characterized by loss of the chromosome 20 p-arm, which harbors the hsa-mir-103–2 previously reported to be downregulated in colorectal tumors^23,24^.

*survClust* is designed to capture the contribution of survival associated molecular features and reduce the influence from those that are not related to the outcome of interest. Figure 4b provides another example that this approach is better in identifying prognostically relevant subtypes compared to the unsupervised clustering approach applied in the original study^25^. *survClust* identified 6 unique CN groups in liver cancer, with significant survival differences among subgroups. The c5 subtype was characterized by high FGA and associated with poor outcome with a median survival time of 0.77 yrs. This cluster was distinguished by a loss of chromosome 15. The c2 subtype was associated with the lowest FGA and a median survival time of 6.81 yrs. The c4 subtype was enriched for *CDKN2A* deletion with a median survival time of 2.15 years. By contrast, unsupervised clustering generated subgroups with distinct molecular differences but did not show any separation in terms of survival.

### Integration of multiple data types enhances the identification of survival distinct subgroups

Figure 5 shows that the integrated *survClust* solution outperformed individual platforms based on the cross validated log rank statistics for multiple cancer types including cervical cancer, head and neck cancer, papillary kidney cancer, lower grade glioma, liver and endometrial cancers. In general, the integrated solutions always emerge at or near the top in performance as compared to the individual platform specific solutions.

**Figure 5:**
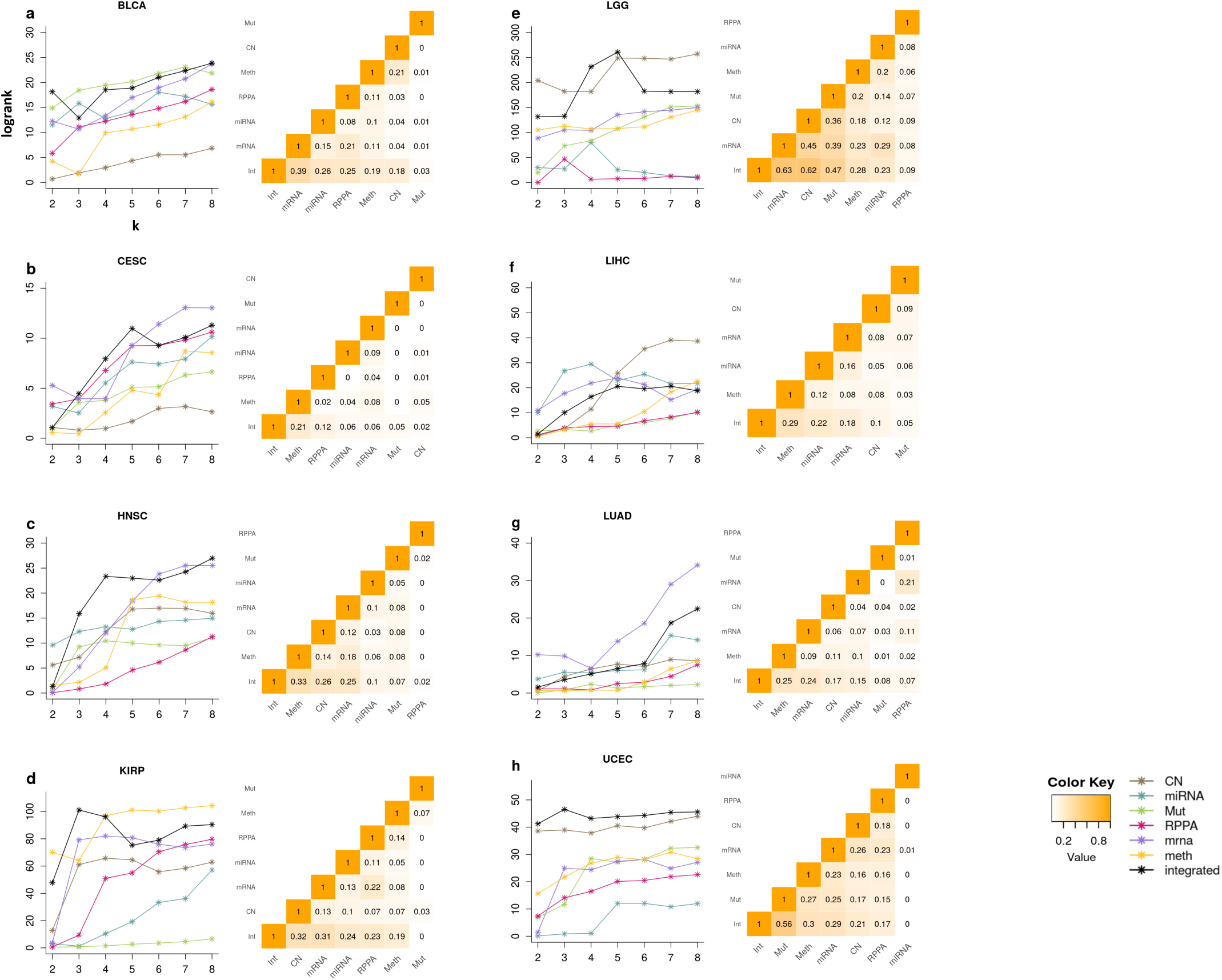
Integration of multiple data types enhances the identification of survival distinct subgroups. **a-h:** Each panel has two plots: the plot on the left summarizes median cross validated log rank statistic across k=2 to 8 (number of clusters). Each line is a data type (see legend), and the black line represents the *survClust* run on integrating all 6 platforms. Plot on the right summarizes the adjusted rand index between cross validated *survClust* solutions of individual data types and the integration of all. In this comparison, the *survClust* solution was chosen for an appropriate k which maximized logrank statistic and minimized the standardized pooled within sum of squares.

Next, we used the adjusted Rand index (RI) to evaluate the concordance between different solutions. RI is calculated as the proportion of sample pairs that are assigned together in the same cluster in one solution versus another, adjusted for what is expected by random chance. It provides an indirect measure of how much a particular data type contributes to the integrated solution. A non-zero adjusted RI across solutions would suggest shared biology across assay types in some tumors. For example, the mutation subtypes of endometrial cancer (**Fig 5h**) have the highest adjusted RI (0.56) as compared to the integrated solution, which is consistent with the fact that POLE and MSI are the two major prognostic subtypes that are predominantly defined through mutation burden (**Fig 3b**). Nevertheless, the integrated solution also clearly shows that there is additional information in DNA methylation, DNA copy number, and mRNA expression being effectively incorporated by *survClust* that improved the survival stratification. In bladder cancer, the integrated solution is most concordant with the mRNA cluster solution (adjusted RI = 0.39), indicating influence by mRNA features towards integration (**Fig 5a**). Classification by mutation data type seemed to have little or no overlap between other assays (adjusted RI close to 0), although the integrated solution retained some information. (adjusted RI=0.03).

The integrated solution classified cervical cancer samples better than rest of the platforms and pointed towards a 5-cluster solution (**Fig 5b**). Interestingly, a high degree of heterogeneity among different platforms was observed as represented by a small adjusted RI across the board. The head and neck cancer integrated solution showed great improvement over individual platforms for k > 2 solutions. The k=4 integrated solution clearly resulted from effective integration of multiple data types including DNA methylation, DNA copy number, and mRNA expression with an adjusted RI of 0.33, 0.26 and 0.25 respectively (**Fig 5c**). In this case, RPPA provided very little information toward the integrated solution.

The integrated survClust analysis stratified papillary kidney cancer type into 3 groups, with CN sharing maximum information with the integrated solution (adjusted RI = 0.32), followed by mRNA (0.31), miRNA (0.24), RPPA (0.23), and Methylation (0.19). Lower grade glioma displayed a wide range of variability among platform type in terms of the logrank statistic (logrank statistic, x-axis from 0-250). The k=5 integrated solution performed the best among the 6 platforms with larger contributions from mRNA (RI = 0.63), copy number (RI = 0.62) and mutation (RI = 0.57) (**Fig 5e**). The integrated solution of liver cancer did not show much improvement over individual assay types. Note that we did not use protein data while integrating as more than half of the samples were not assayed with the protein platform (RPPA, n=182; integrated n=371). miRNA, mRNA and copy number showed high median logrank statistics over rounds of cross-validation demonstrating their role as potential prognostic classifiers.

## DISCUSSION

We proposed a supervised clustering algorithm, *survClust*, that directly incorporates time to event (e.g., death, disease progression) information with molecular features to stratify patients into clinically relevant subtypes. We further developed two visualization tools, *circomap* and *panelmap* for displaying and annotating the resulting stratification. As more clinically annotated genomic data is becomes available as a result of clinical sequencing programs^26,27^, our method will provide a useful tool to facilitate patient stratification for clinical decision making. In this study, we analyzed 18 cancer types in ∼ 6200 tumors. Each disease type was classified by *survClust* based on six molecular assays – somatic point mutation, DNA copy number, DNA methylation, mRNA expression, miRNA expression, protein expression and integration of the aforementioned six assays.

The supervised clustering approach provides a more direct way to identify survival associated molecular subclasses, often leading to substantially more distinct survival subgroups than those existing molecular subclasses obtained by unsupervised clustering. For example. The integrated *survClust* stratification of the hepatocellular carcinomas (LIHC) was associated with a survival log-rank statistic of 45.19 (P<0.001) versus 1.69 (P=0.42) under the unsupervised clustering solution (**Supplementary Table 2, Supplementary Fig 8**), suggesting that *survClust* is a more powerful approach for identifying outcome-associated subtypes. Supplementary Tables 2-7 show comparisons of the log-rank statistics in survival differences across the various integrated and individual platform *survClust* solutions with those from existing molecular clustering solutions reported in the TCGA publications (wherever available). Note that survClust solutions have all been cross-validated to avoid overfitting.

The outcome-weighted learning framework we propose in this study can be extended to model binary outcome types such as treatment response or toxicity (which is an important outcome category in immunotherapy settings). In addition, the integration framework can facilitate the inclusion of other data modalities including histopathological data and radiological images.

## METHODS

### survClust workflow

Let ***X***_***m***_ be the *m*^*th*^ (m=1,…,M) data type of dimension *N*_*m*_ (number of samples in *m*^*th*^ data type, can vary) by *p*_*m*_ (number of features). Data types may consist of continuous (gene expression, copy number log-ratio, DNA methylation, miRNA, protein expression) or binary (mutation status) data. Overall survival is defined as time from diagnosis to death or last follow-up. The data needs to be pre-processed as described in **Supplementary Information.**

For a pair of two samples *a* and *b*, **the weighted distance**^7^ is calculated as follows:

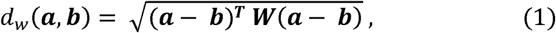

where, ***a*** and ***b*** bare feature vectors of length *p*, for samples *a* and *b* respectively, ***W*** is a *p* × *p* diagonal weight matrix with ***W*** = *diag* {*w*_l_, …,*w*_*p*_}. Samples are close to each other when the value of *d*_*w*_ is small and dissimilar when *d*_*w*_ is large.

The weights *w*_*j*_ (*j* = 1,…,*p*) are obtained by fitting a univariate cox proportional hazards model for each feature:

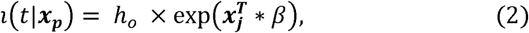

where, represents the survival time, ***x***_***j***_ is the *j*^*th*^ column of matrix ***X*** of length *N, h*_0_, is the baseline hazard function, *β* is the regression coefficient and exp(*β*) is the Hazard Ratio (HR).

We consider the absolute value of HR on the logarithmic scale as the weight w. An HR=1 corresponds to the null that the feature is not associated with survival. This is reflected in a log(1) =0 weighting in the distance matrix. Since w is a diagonal matrix with diagonal element *w*_*j*_ (*j* = 1,…,*p*), we can simply use euclidean distance for computing distances if we transform the data as follows:

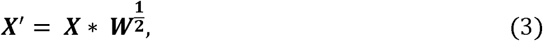

Euclidean distances are sensitive to scale of the observations. After incorporating weights, we standardize the data by its grand total:

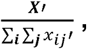

where, ∑_***i***_ ∑_***j***_ *x*_*ij*_ ′ is the grand total of weighted matrix ***X***′, with *i* rows (N samples) and *j* columns (p features). Then, one can compute the pairwise distance between samples *a* (*i*= 1) and *b*(*i* = 2) as:

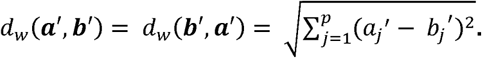

Conversely, a weighted distance matrix ***D*** is calculated for all pairwise samples across *M* data types. All samples having full survival information are kept, and the union of all samples (*N*_*union*_) across *M* data types is utilized when analyzing a wide number of samples. Non-overlapping samples in data types are added as *NA* to have a uniform set of *N*_*union*_ samples.

The integrated weighted distance matrix is calculated by averaging over the weighted distance matrices:

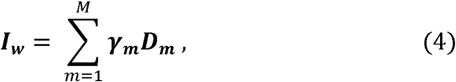

where 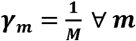. The integrated weighted matrix ***I***_***w***_, averages the inter-and intra-sample similarity profiles over the *M* data types. ***I***_***w***_ is then processed by *survClust* via classical multidimensional scaling (MDS) ^28^ and clustered using k-means^29^. Classical MDS assumes Euclidean distances; however, in cases of non-Euclidean distances, Mardia et al^30^ provided a method to obtain the resulting positive semidefinite scalar product matrix. Note that *I*_*w*_ follows the Euclidean norm and hence can be represented in *n* − 1 dimensions. The strong assumption of the Euclidean norm is also important for k-means, as it is essentially trying to assign samples to the closest centroid or calculating the sum of squared deviations from centroids.

### Weighted distance metric for mutation data

Somatic mutation data is represented as a binary data matrix where each entry is coded as 1 if the *j*^th^ gene is mutated in the *i*^th^ sample, and 0 otherwise. A challenge with the mutation data matrix is the sparsity. It is known that somatic mutation data exhibit a long-tailed distribution in which a relatively small number of variants appear in tumors frequently while the vast majority of variants occur extremely infrequently. We consider genes that are mutated in > 1% of the sample. After incorporating weights, this data is no longer binary, but it still remains sparse. Due to such data sparsity, computing Euclidean distance is not appropriate and may lead to inflated distance measures^31^. To combat this problem, we propose a weighted binary distance metric for such a scenario as described below.

Let 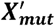 be the weighted mutation data matrix (see Equation. 3) of dimension *N* (samples) by *p* (genes). Then, the pairwise distance between sample vectors ***a*** and ***b*** is calculated as follows:

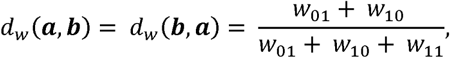

where

*w*_0l_ = sum of weights of *p* features that are zero in sample vector ***a*** but non-zero in sample vector ***b*;**

*w*_l0_ = sum of weights of *p* features that are non-zero in sample vector ***a*** but zero in sample vector ***b*;**

*w*_ll_ = sum of weights of *p* features that are non-zero in sample vector ***a*** and non-zero in sample vector ***b*.**

Note that, *d*_*w*_ (***a, b***) is a proportion of sum of effect sizes in which only one is non-zero amongst those in which at least one is non-zero. ^32^

### Cross-validation

*survClust* classifies sample populations by incorporating outcome information. Resulting clusters are overly optimistic and need to be cross validated to arrive at more generalizable solutions. The *cv.survclust* function provides cross validation for the desired number of folds and outputs cross-validated solution labels. In the results shown above, we performed 5-fold cross validation as follows: (1) Split the data into 5 random partitions, label 4 of them as the training sets and the remaining one as the test set. (2) The weighted distance matrix was calculated from the training data set alone (Eq.1). *survClust* clustering was performed to arrive at outcome weighted labels in the training set. (3) test labels were predicted according to training labels (4) Step 2 was repeated until predictions were made on all 5 test data sets across all 5 folds. (6) clusters were tracked by *centroid relabeling* (**Supplementary Note 1.3**) across folds, and we obtained outcome weighted class labels for our entire dataset. This concluded one round of cross-validation. All results shown here are results from cross-validated labels across 50 rounds of cross-validation. Cluster meaning was preserved across rounds of cross validation via a similar approach to centroid relabeling. The final label for a sample was assigned to a class to which it was predicted in the maximum number of rounds. This is achieved by another function called *consensus.summary*.

### Choice of the number of clusters k

The logrank test statistic and standardized pooled within-cluster sum of squares were calculated from cross-validated labels to choose an appropriate,.

### Logrank test statistic

For a particular *k* cluster solution we have *k* cross-validated labels. Each class is distinct in survival and we can compare the difference between classes using the logrank test statistic as follows^33^:

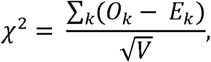

where, *O*_*k*_ = observed number of events in the *k*^*th*^ group over time, *E*_*k*_ = expected number of events in the *k*^*th*^ group over time and *V* = ∑*Var* (*O*_*k*_ − *E*_*k*_) = ∑*V*_*k*_.

### Standardized pooled within-cluster sum of squares

Here we calculate the pooled within-cluster sum of squares and standardize it by the total sum of squares similar to methodology used in the gap statistic^34^ to select the appropriate number of clusters. Suppose that the final labels have clustered the data into *k* clusters *C*_l_, *C*_2_, …. *C*_*k*_, with *C*_*r*_ denoting the indices of observations in cluster *r*, and *n*_*r*_ = |*C*_*r*_ |. Let

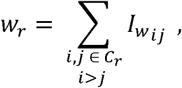

where *w*_*r*_ is the sum of all pairwise distances in cluster *r*, {*ij*} represents a pair of samples belonging to a cluster *C*_*r*_ and ***I***_***w***_ is calculated from Eq 4. Then the standardized pooled within-cluster sum of squares is calculated as:

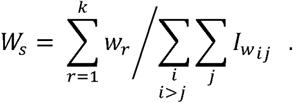

Here *W*_*s*_ decreases monotonically as the number of clusters *k* increases. The optimal number of clusters is where *W*_*s*_ is minimized and creates an ‘elbow’ or a point of inflection, where addition of more clusters does not improve cluster separation. Another property of *W*_*s*_ is that it can be used to compare amongst different datasets as it lies between 0 and 1 after standardization. This is useful in comparing *survClust* runs between individual data types and when we integrate them.

### Simulation

Continuing from the simulation study presented in **Fig 1**, we go into detail about cross-validation and how to chose **k** for a *survClust* run. In **Fig 1**, the input matrix was subjected to 50 rounds of 3-fold cross-validation (2/3 training and 1/3 test. The *survClust* fit for a cluster ***k*** based on training data from each fold was used to predict cluster membership for the remaining 1/3 test data. Final sample labels were aggregated over all folds and cluster meaning was preserved across folds via centroid relabeling. (**See Supplementary Note 1.3**).

Logrank test statistic and standardized pooled within-cluster sum of squares was calculated for the consolidated test labels over 3-folds for each round. **Supplementary Fig 1(c)** summarizes these metrics for 50 rounds of cross validation for k=2-7. We see that logrank is maximized for k=3, and the standardized pooled within-cluster sum of squares elbows at k=3, pointing to the optimal selection of k at k=3. The final class labels are assigned by consolidating solutions across all folds in all rounds of cross validations.

### Implementation and availability

*survClust* is freely available as an R package at (https://github.com/arorarshi/survClust).

For k-means clustering, we used the k-means implementation in the R base package. For multidimensional scaling, we used the *cmdscale* function in base R. The weighted distance metric for binary data was programmed in C++ with R extension using *Rcpp* package, which is computationally fast. Hazard ratios were derived from the cox proportional hazard model came from the R *survival* package. Kaplan Meier curves were plotted using *ggsurvplot* in package *survminer*. Beeswarm plots were made using R package *beeswarm*. Mutation data along with relevant clinical annotations were plotted using *panelmap* (https://github.com/arorarshi/panelmap). The *circlize* R package was used to make pan-cancer plots and the code used to plot these is available in a function called *circomap* https://github.com/arorarshi/panelmap#example---circomap)

Below is the workflow of proposed *survClust* method:

1. **getDist** – Compute a weighted distance matrix across given *m* data types. Standardization and accounting for non-overlapping samples is also accomplished in this step.
2. **combineDist** – Integrate *m* data types by averaging over *m* weighted distance matrices.
3. **survClust** and **cv.survclust** – Estimate the survClust solution for a given cluster number ***k*** based on the weighted and integrated distance matrix. Optimal k is estimated via cross-validation. Use the chosen k and the cross-validated results to arrive at final class labels. Cross-validated results are assessed over the following performance metrics – the logrank statistic, standardized pooled within-cluster sum of squares and cluster solutions with class size less than 5 samples.

## DECLARATIONS

### Availability of Data and materials

The *survClust* algorithm’s software implementation is available at https://github.com/arorarshi/survClust. Simulated dataset and simulated survival dataset are also available on GitHub. All genomics and clinical data was downloaded from https://portal.gdc.cancer.gov/. *panelmap* and *circomap* are available at https://github.com/arorarshi/panelmap

### Author’s contributions

A.A. A.B.O., V.E.S. and R.S. designed the research. A.A. made software implementations and analyzed the data. A.A. A.B.O., V.E.S. and R.S. wrote the paper.

## Acknowledgements

The research was supported by the National Cancer Institute, Award CA008748.

## REFERENCES

1 Shen, R. L., Olshen, A. B. & Ladanyi, M. Integrative clustering of multiple genomic data types using a joint latent variable model with application to breast and lung cancer subtype analysis. Bioinformatics 25, 2906–2912, doi:10.1093/bioinformatics/btp543 (2009).

2 Mo, Q. et al. Pattern discovery and cancer gene identification in integrated cancer genomic data. Proc Natl Acad Sci U S A 110, 4245–4250, doi:10.1073/pnas.1208949110 (2013).

3 Vaske, C. J. et al. Inference of patient-specific pathway activities from multi-dimensional cancer genomics data using PARADIGM. Bioinformatics 26, i237–i245, doi:10.1093/bioinformatics/btq182 (2010).

4 Hoadley, K. A. et al. Multiplatform Analysis of 12 Cancer Types Reveals Molecular Classification within and across Tissues of Origin. Cell 158, 929–944, doi:10.1016/j.cell.2014.06.049 (2014).

5 Wang, B. et al. Similarity network fusion for aggregating data types on a genomic scale. Nat Methods 11, 333–337, doi:10.1038/nmeth.2810 (2014).

6 Ramazzotti, D., Lal, A., Wang, B., Batzoglou, S. & Sidow, A. Multi-omic tumor data reveal diversity of molecular mechanisms that correlate with survival. Nature Communications 9, 4453, doi:ARTN 4453 10.1038/s41467-018-06921-8 (2018).

7 Xing, E. P., Jordan, M. I., Russell, S. J. & Ng, A. Y. in Advances in neural information processing systems. 521–528.

8 Hoadley, K. A. et al. Cell-of-Origin Patterns Dominate the Molecular Classification of 10,000 Tumors from 33 Types of Cancer. Cell 173, 291–304 e296, doi:10.1016/j.cell.2018.03.022 (2018).

9 Thorsson, V. et al. The Immune Landscape of Cancer. Immunity 48, 812–830 e814, doi:10.1016/j.immuni.2018.03.023 (2018).

10 Malta, T. M. et al. Machine Learning Identifies Stemness Features Associated with Oncogenic Dedifferentiation. Cell 173, 338–354 e315, doi:10.1016/j.cell.2018.03.034 (2018).

11 Sanchez-Vega, F. et al. Oncogenic Signaling Pathways in The Cancer Genome Atlas. Cell 173, 321–337 e310, doi:10.1016/j.cell.2018.03.035 (2018).

12 Novembre, J. et al. Genes mirror geography within Europe. Nature 456, 98–U95, doi:10.1038/nature07331 (2008).

13 Network, C. G. A. R. Comprehensive, integrative genomic analysis of diffuse lower-grade gliomas. New England Journal of Medicine 372, 2481–2498 (2015).

14 Yan, H. et al. IDH1 and IDH2 mutations in gliomas. N Engl J Med 360, 765–773, doi:10.1056/NEJMoa0808710 (2009).

15 Network, C. G. A. R. Comprehensive molecular characterization of papillary renal-cell carcinoma. New England Journal of Medicine 374, 135–145 (2016).

16 Alexandrov, L. B. et al. Signatures of mutational processes in human cancer. 500, 415 (2013).

17 Zhou, Q. et al. Worldwide research trends on aristolochic acids (1957–2017): Suggestions for researchers. 14, e0216135 (2019).

18 Getz, G. et al. Integrated genomic characterization of endometrial carcinoma. Nature 497, 67–73, doi:10.1038/nature12113 (2013).

19 Newman, A. M. et al. Robust enumeration of cell subsets from tissue expression profiles. Nat Methods 12, 453–457, doi:10.1038/nmeth.3337 (2015).

20 Chae, Y. K. et al. Mutations in DNA repair genes are associated with increased neo-antigen load and activated T cell infiltration in lung adenocarcinoma. Oncotarget 9, 7949–7960, doi:10.18632/oncotarget.23742 (2018).

21 Olshen, A. B., Venkatraman, E. S., Lucito, R. & Wigler, M. Circular binary segmentation for the analysis of array-based DNA copy number data. Biostatistics 5, 557–572, doi:10.1093/biostatistics/kxh008 (2004).

22 Bellacosa, A. et al. Molecular Alterations of the Akt2 Oncogene in Ovarian and Breast Carcinomas. International Journal of Cancer 64, 280–285, doi:DOI 10.1002/ijc.2910640412 (1995).

23 Sheffer, M. et al. Association of survival and disease progression with chromosomal instability: a genomic exploration of colorectal cancer. Proc Natl Acad Sci U S A 106, 7131–7136, doi:10.1073/pnas.0902232106 (2009).

24 Cummins, J. M. et al. The colorectal microRNAome. Proc Natl Acad Sci U S A 103, 3687–3692, doi:10.1073/pnas.0511155103 (2006).

25 Ally, A. et al. Comprehensive and integrative genomic characterization of hepatocellular carcinoma. 169, 1327–1341. e1323 (2017).

26 Zehir, A. et al. Mutational landscape of metastatic cancer revealed from prospective clinical sequencing of 10,000 patients. 23, 703 (2017).

27 Micheel, C. M. et al. American association for cancer research project genomics evidence neoplasia information exchange: from inception to first data release and beyond—lessons learned and member institutions’ perspectives. 2, 1–14 (2018).

28 Torgerson, W. S. Theory and methods of scaling. (1958).

29 Hartigan, J. A. & Wong, M. A. Algorithm AS 136: A k-means clustering algorithm. J Journal of the Royal Statistical Society. Series C 28, 100–108 (1979).

30 Mardia, K. V. & Methods. Some properties of clasical multi-dimesional scaling. J Communications in Statistics-Theory 7, 1233–1241 (1978).

31 Legendre, P. & Gallagher, E. D. Ecologically meaningful transformations for ordination of species data. Oecologia 129, 271–280, doi:10.1007/s004420100716 (2001).

32 Martin, N. & Maes, H. Multivariate analysis. (Academic press London, 1979).

33 Harrington, D. P. & Fleming, T. R. A Class of Rank Test Procedures for Censored Survival-Data. Biometrika 69, 553–566, doi:DOI 10.1093/biomet/69.3.553 (1982).

34 Tibshirani, R., Walther, G. & Hastie, T. Estimating the number of clusters in a data set via the gap statistic. Journal of the Royal Statistical Society Series B-Statistical Methodology 63, 411–423, doi:Doi 10.1111/1467-9868.00293 (2001).

