## Supplementary Notes for "Pan-cancer identification of clinically relevant genomic subtypes using outcome-weighted integrative clustering"

### 1. Data Pre-Processing

Data types may consist of continuous (gene expression, copy number log-ratio, DNA methylation, miRNA, protein expression) or binary (mutation status) data. Each data type is pre-processed by normalization and standardized as follows –

$$\boldsymbol{X}_{\boldsymbol{standardized}} = \frac{\boldsymbol{X} - \overline{\boldsymbol{X}}}{\boldsymbol{sd}(\boldsymbol{X})}$$

where X is a data type.

Copy Number data was segmented using CBS[1] and reduced to non-redundant regions of alterations using the CNregion function in iClusterPlus[2] with default epsilon of 0.001, keeping in mind that the total numbers of features don't exceed 10,000. For DNA methylation, mRNA expression and miRNA expression, if a certain feature had more than 20% missing data, that feature was removed and remaining were used for analysis. For mRNA expression, we further removed genes having a mean expression lower than the threshold of mean expression of lower 10% quantile. Similarly, methylation probes with mean beta values < 0.1 and > 0.9 were discarded. Genes harboring mutants in less than 1% of the samples were removed. Missing data was imputed using KNN imputation[3]. Data for each cancer type and assay was extracted from pan-cancer study[4]. A summary of features and samples across all cancer types analyzed is shown in Supplementary Table 1. Mutation signature analysis was run on the resulting maf file using (https://github.com/mskcc/mutation-signatures)[5]. Blood biomarker data was derived from CIBERSORT[6].

### 2. Running survClust

*survClust* was run on all cancer types reported in Supplementary Table 1. Results are summarized according to each cancer type. Each cancer type was run for each of the 6 platforms - somatic mutation, DNA copy number, DNA methylation, mRNA expression, miRNA expression and RPPA data, and integrating all six.
Final results were compiled after running *cv.survclust* for 5 folds for 50 rounds of cross validation. Optimal $k$ was chosen after assessing cross-validated logrank statistic and standardized pooled within cluster sum of squares. Overfitting was avoided by omitting cluster solution with less than 5 samples. Note that, one can considerably reduce the time to run survClust by reducing the feature space.

### 3. Simulation details

We present here two examples of simulations to go over the conceptualization of survClust.

#### 3.1 Simulation 1

We simulated a data matrix with 300 samples and 300 features. Out of which 15 features, $f$, were coded for survival relevant information for 100 samples each in cluster1,2 and 3 respectively -

$$\boldsymbol{C1}_{(\boldsymbol{f=15})} \sim N(0,1),\ \ \boldsymbol{C2}_{(\boldsymbol{f=15})} \sim N(-1.5,1), \boldsymbol{C3}_{(\boldsymbol{f=15})} \sim N(1.5,1) \qquad (\mathbf{1})$$

Next 15 were coded for survival unrelated features and formed the competing cluster. Survival association was perturbed by permuting the samples, which dropped the associated survival information within features but retained the molecular distinction between clusters.

$$\boldsymbol{C1}_{(\boldsymbol{f_{permute}=15})} \sim N(0,1),\ \ \boldsymbol{C2}_{(\boldsymbol{f_{permute}=15})} \sim N(-1.5,1),\ \boldsymbol{C3}_{(\boldsymbol{f_{permute}=15})} \sim N(1.5,1)$$

$$(\mathbf{1.1})$$

And remaining 270 features were simulated as noise –

$$\boldsymbol{C1}_{(\boldsymbol{f=270})} \sim N(0,1),\ \ \boldsymbol{C2}_{(\boldsymbol{f=270})} \sim N(0,1), \boldsymbol{C3}_{(\boldsymbol{f=270})} \sim N(0,1) \qquad (\mathbf{1})$$

Survival correlation was imposed by simulating C1, C2 and C3, to have a median survival time of 4, 3 and 2 years respectively. The simulated survival data observed right censoring, with each individual having failure time as T and censoring time as C, and time-to-event data as follows –

$$Y = \min(T, C) = T \perp C$$
$$\delta = \begin{cases} 1, if\ T = Y \\ 0\ \ otherwise \end{cases}$$
$$T \sim \exp(\lambda)\ , C \sim U(a, b) \qquad (\mathbf{2})$$

Where, λ=median survival rate, such as, a median survival rate of 4 years is - log(2)/4. $a$,$b$ are minimum and maximum follow up times.
We created two such data types to showcase integration capabilities of survClust (**Supplementary Fig 1**). We ran survClust and cross-validated runs for k=2-7 for 50 rounds. Optimum $k$ is chosen where logrank is maximized and standardized pooled within-cluster sum of squares is minimized. Results are shown in Figure 1.

#### 3.2 Simulation 2

We show another simulation example, where we simulate two data types where the truth is a 4-class solution shown in **Supplementary Figure 2**, such that:

$$\textbf{Data type 1}:\ \boldsymbol{C1}_{(\boldsymbol{f=15,N=100})} \sim N(0,1),\ \ \boldsymbol{C2}_{(\boldsymbol{f=15,N=100})} \sim N(-1.5,1),$$

$$\boldsymbol{C3}_{(\boldsymbol{f=15,N=250})} \sim N(1.5,1)$$

$$\textbf{Data type 2}:\ \boldsymbol{C1}_{(\boldsymbol{f=15,N=100})} \sim N(1.5,1),\ \ \boldsymbol{C2}_{(\boldsymbol{f=15,N=100})} \sim N(0.5,1),$$

$$\boldsymbol{C3}_{(\boldsymbol{f=15,N=100})} \sim N(0,1), \boldsymbol{C4}_{(\boldsymbol{f=15,N=150})} \sim N(-0.5,1)$$

Data type 1 shows strong molecular association, whereas data type 2 shows strong survival association and weak molecular association.

Survival correlation was imposed by simulating C1, C2, C3 and C4 with median survival time of 5, 4, 3 and 2 years respectively. See (2) and Supplementary Note 1.1. The goal here was to see if survClust can identify 4 distinct survival groups when a data type has a strong 3-class molecular structure (as shown in Data type1). Results shown in Supplementary Figure 2.

### 4. Centroid Re-labelling

We performed cross validation to avoid over fitting and to arrive at coherent survClust results. Each fold predicts test labels according to its training set, and at the end of cross validation we have prediction of each sample to a class. However, the class labels are meaningless across folds and one needs to be careful when aggregating labels to get a full solution. We define, centroid relabeling method to solve this problem.

Let $N$ be total number of samples to classify. We perform a $F$ fold cross validation, where each fold predicts a test set, such that,

$\sum_{f=1}^{F} test\ set_f = N.$

Assume we have the following class label prediction for test set for $k$-class solutions across $F$ folds.

$test\ set_1 = a_{11}, a_{12}, a_{13}, a_{24}, a_{35}, a_{26}..a_{ki};$; $test\ set_2 = a_{11}, a_{12}, a_{13}, a_{24}, a_{35}, a_{26}..a_{ki};$ $\ldots test\ set_f = a_{11}, a_{12}, a_{13}, a_{24}, a_{35}, a_{26}..a_{ki}$

Where $a_{ki}$= $i^{th}$ sample belonging to $k^{th}$ class in a test set fold $f$. One can clearly see that $k$ labels across $f$ folds are meaningless and simply group alike samples together and unlike samples separately in different clusters. To classify all $N$ samples, we need to consolidate the labels across $F$ folds.

Lets consider the following example where we know the true class of each sample -

| **Sample** | **Class** |
|---|---|
| sample1 | 1 |
| sample2 | 2 |
| sample3 | 3 |
| sample4 | 1 |
| sample5 | 2 |
| sample6 | 3 |
| sample7 | 1 |
| sample8 | 2 |
| sample9 | 3 |

Now we perform 3-fold cross validation, using 2/3rd of the samples (n=6) to predict the remaining 1/3rd (n=3) samples in each fold.

| test set prediction- Fold 1 | | Fold 2 | | Fold 3 | |
|---|---|---|---|---|---|
| sample4 | class1 | sample5 | class1 | sample6 | class1 |
| sample2 | class2 | sample3 | class2 | sample1 | class2 |
| sample9 | class3 | sample7 | class3 | sample8 | class3 |

As seen, these class labels are meaningless, but they group alike samples together and unlike samples separately. For example, Sample4, Sample5 and Sample6 are grouped in class1 across folds, but we know that their true label is class1, class2 and class3 respectively. We arrive at consolidated labels as follows-

Step 1- Calculate centroids of each $k (k = 3)$ class in each fold $f (f = 3)$.

$$Fold\ 1\ centroids - \ o_{11}, o_{21}, o_{31}$$

$$Fold\ 2\ centroids - \ o_{12}, o_{22}, o_{32}$$

$$Fold\ 3\ centroids - \ o_{13}, o_{23}, o_{33}$$

Where, $o_{kf}; k = k\ class$ and $f = f\ fold$

Where, $o_{kf} = \frac{\sum_k a_{k1}, + a_{k2} + a_{k3} + \cdots + a_{ki}}{i}$; centroid calculation for $f^{th}$ fold and $k^{th}$ class and $i$ samples belonging to that $k$ class.

Step 2- Run kmeans on the centroids vectors

$$kmeans( o_{11}, o_{12}, o_{13}, o_{21}, o_{22}, o_{23}, o_{31}, o_{32}, o_{33})$$

Step 3 – Get kmeans class labels on the centroids, and relabel each sample accordingly.

| Kmeans output | Centroids |
|---|---|
| k=1 | $o_{11}, o_{32}, o_{23}$ |
| k=2 | $o_{21}, o_{12}, o_{33}$ |
| k=3 | $o_{31}, o_{22}, o_{13}$ |

Re-assigned class labels -

| Fold1 | | Fold2 | | Fold3 | |
|---|---|---|---|---|---|
| Sample4 | class1 | Sample5 | Class2 | Sample6 | Class3 |
| Sample2 | class2 | Sample3 | Class3 | Sample1 | Class1 |
| Sample9 | class3 | Sample7 | Class1 | Sample8 | Class2 |

This same strategy is used to relabel class membership of samples across rounds of cross validation, where each round has predicted class labels after each round of cross validation.

For a particular $k$ each round $r$ $(1,2,3 \ldots, R)$ of cross validation, we have relabeled class labels as described above, as $cv\ labels_1, cv\ labels_2, \ldots\ cv\ labels_R$.

Consolidate labels across all $R$ rounds to arrive at the final solution as follows.

Compute centroids for each $k$ class in each round $R$ (same as Step1).

$r_1\, centroids -\ o_{11}, o_{21}, o_{31}, \ldots\ o_{k1}; r_2\ centroids -\ o_{12}, o_{22}, o_{32}, \ldots\ o_{k2}; r_R\ centroids -\ o_{1R}, o_{2R}, o_{3R}, \ldots\ o_{kR}$

Proceed to Step 2 and Step 3 to get relabeled class labels across each $r$ round. Now the cluster membership has same meaning across all rounds of cross validation. A final class label is picked that is seen maximum number of times over $R$ rounds.

### References


1. Olshen, A. B., Venkatraman, E., Lucito, R. & Wigler, M. J. B. Circular binary segmentation for the analysis of array-based DNA copy number data. 5, 557-572 (2004).

2. Mo, Qianxing, and Ronglai Shen.iClusterPlus: integrative clustering of multiple genomic data sets.(2013).

3. Trevor Hastie, Robert Tibshirani, Balasubramanian Narasimhan and Gilbert Chu (2017). impute: impute: Imputation for microarray data. Rpackage version 1.50.1.

4. Hoadley, K. A. *et al.* Cell-of-origin patterns dominate the molecular classification of 10,000 tumors from 33 types of cancer. 173, 291-304. e296 (2018).

5. Alexandrov, L. B. *et al.* Signatures of mutational processes in human cancer. 500, 415 (2013)

6. Newman, A. M. *et al.* Robust enumeration of cell subsets from tissue expression profiles. 12, 453 (2015).
