## Supplementary Figures and Tables for "Pan-cancer identification of clinically relevant genomic subtypes using outcome-weighted integrative clustering"

| **Content** | | **Page Number** |
|---|---|---|
| **Supplementary Table 1** | Summarizing input data of 18 cancer types across number of samples analyzed and total number of features that went into clustering for mutation, Copy Number (CN), Methylation, mRNA expression, miRNA and Protein | 2 |
| **Supplementary Table 2** | Comparison of *survClust* **integrated** solution versus unsupervised clustering results from published TCGA studies | 3 |
| **Supplementary Table 3** | summary of *survClust* **Copy Number** solution vs unsupervised TCGA solutions on logrank | 4 |
| **Supplementary Table 4** | summary of *survClust* **Methylation expression** solution vs available TCGA solutions on logrank | 5 |
| **Supplementary Table 5** | summary of *survClust* **mRNA expression** solution vs available TCGA solutions on logrank | 6 |
| **Supplementary Table 6** | summary of *survClust* **miRNA expression** solution vs available TCGA solutions on logrank | 7 |
| **Supplementary Table 7** | summary of *survClust* **Protein expression** solution vs available TCGA solutions on logrank | 8 |
| **Supplementary Table 8** | Cross tabulation of OV copy number *survClust* labels vs Copy number change in *AKT* gene | 9 |
| **Supplementary Figure 1** | Understanding survClust simulation and analysis | 10 |
| **Supplementary Figure 2** | Another simulation example | 11 |
| **Supplementary Figure 3** | *survClust* integrated solution of LGG | 12 |
| **Supplementary Figure 4** | *survClust* identifies TMB patterns across cancer types | 13 |
| **Supplementary Figure 5** | CD8 T-cell distribution of *survClust* mutation classes of various cancer types | 14 |
| **Supplementary Figure 6** | *survClust* identifies FGA patterns across cancer types, Global view of Copy Number Change | 15 |
| **Supplementary Figure 7** | *survClust* identifies FGA patterns across cancer types, Kaplan Meier curves | 16 |
| **Supplementary Figure 8** | *survClust* integrated solution of various cancer types | 17 |

**Supplementary Table 1** – Summarizing input data of 18 cancer types across number of samples analyzed and total number of features that went into clustering for mutation, Copy Number (CN), Methylation, mRNA expression, miRNA and Protein

| cancer type | Total samples analyzed | mutation | CN | Methylation | mRNA | miRNA | RPPA |
|---|---|---|---|---|---|---|---|
| **BLCA** | 411 | 8867 | 3353 | 9293 | 15695 | 553 | 189 |
| **CESC** | 293 | 5579 | 8471 | 9743 | 15771 | 565 | 193 |
| **COAD** | 432 | 7064 | 1909 | 9432 | 15331 | 494 | NA |
| **ESCA** | 185 | 1790 | 6508 | 9938 | 16997 | 535 | 193 |
| **HNSC** | 528 | 6304 | 7942 | 9662 | 15840 | 573 | 191 |
| **KIRP** | 289 | 1506 | 2274 | 7697 | 15675 | 502 | 190 |
| **LGG** | 512 | 665 | 4383 | 8866 | 15920 | 592 | 190 |
| **LIHC** | 372 | 2490 | 3719 | 9900 | 15106 | 550 | 190 |
| **LUAD** | 515 | 4803 | 2053 | 11546 | 15979 | 542 | 189 |
| **LUSC** | 503 | 4485 | 3650 | 9019 | 16227 | 543 | 189 |
| **MESO** | 86 | 43 | 3416 | 8506 | 15798 | 558 | 190 |
| **OV** | 537 | 2148 | 2432 | 7175 | 16847 | 531 | 189 |
| **PAAD** | 184 | 3494 | 4521 | 9061 | 16193 | 565 | 190 |
| **SARC** | 261 | 5868 | 4024 | 8840 | 15574 | 463 | 193 |
| **STAD** | 423 | 5595 | 4180 | 10583 | 15416 | 510 | 193 |
| **UCEC** | 542 | 9175 | 2320 | 8650 | 16811 | 540 | 189 |
| **UCS** | 56 | 1149 | 3706 | 8288 | 16194 | 633 | 193 |
| **UVM** | 80 | 87 | 749 | 6595 | 14626 | 557 | NA |

**Supplementary Table 2** – Comparison of *survClust* **integrated** solution versus unsupervised clustering results from published TCGA studies. The logrank test statistics (cross-validated) for survival association of the subtypes are reported.

| Cancer type | survClust | Unsupervised clustering | |
|---|---|---|---|
| | Log-rank statistic | Log-rank | Algorithm |
| Bladder | 22.65 | 4.8 | COCA |
| Cervical | 12.47 | -- | -- |
| Colorectal | 5.2 | 0.61 | PARADIGM |
| Esophageal | 1.73 | 1.08 | iCluster |
| Head and neck | 24.41 | 8.73 | Manual |
| Kidney papillary | 129.79 | 79.09 | COCA |
| Low grade glioma | 288.68 | 255.65 | COCA |
| Liver | 45.19 | 1.69 | iCluster |
| Lung adeno | 6.46 | 9.09 | iCluster |
| Lung squamous | 5.06 | 0.81 | iCluster |
| Mesothelioma | 7.86 | -- | -- |
| Ovarian | 7.34 | 4.93 | |
| Pancreas | 14.7 | -- | -- |
| Soft tissue sarcoma | 17.94 | -- | -- |
| Stomach | 5.2 | 2.65 | Manual |
| Endometrial | 45.65 | 35.65 | Manual |
| Uterine | 0.92 | 0.71 | |
| Uveal melanoma | 16.16 | -- | -- |

**Supplementary Table 3** – summary of *survClust* **Copy Number** solution vs unsupervised TCGA solutions on logrank (columns 1 and 2). Total number of samples analyzed via *survClust* (column 3), and total samples analyzed via unsupervised clustering (column 4)

| Cancer type | survClust logrank | unsup logrank | survClust (N) | unsup(N) |
|---|---|---|---|---|
| **BLCA** | 3.45 | NA | 405 | NA |
| **CESC** | 0.76 | NA | 280 | NA |
| **COAD** | 5.9 | NA | 409 | NA |
| **ESCA** | 0.96 | NA | 182 | NA |
| **HNSC** | 18.64 | 9.86 | 517 | 279 |
| **KIRP** | 57.73 | 46.93 | 281 | 161 |
| **LGG** | 246.74 | NA | 507 | NA |
| **LIHC** | 51.95 | 3.16 | 362 | 193 |
| **LUAD** | 5.96 | NA | 496 | NA |
| **LUSC** | 3.1 | NA | 486 | NA |
| **MESO** | 3.45 | NA | 86 | NA |
| **OV** | 27.13 | NA | 558 | NA |
| **PAAD** | 8.96 | NA | 182 | NA |
| **READ** | 3.92 | NA | 148 | NA |
| **SARC** | 12.59 | NA | 252 | NA |
| **STAD** | 2.18 | NA | 419 | NA |
| **UCEC** | 35.54 | 19.64 | 518 | 537 |
| **UCS** | NA | NA | NA | NA |
| **UVM** | 16.53 | NA | 80 | NA |

**Supplementary Table 4** – summary of *survClust* **Methylation expression** solution vs available TCGA solutions on logrank (columns 1 and 2). Total number of samples analyzed via *survClust* (column 3), and total samples analyzed via TCGA (column 4)

| Cancer type | survClust logrank | TCGA logrank | survClust(N) | TCGA(N) |
|---|---|---|---|---|
| **BLCA** | 9.89 | NA | 411 | NA |
| **CESC** | 3.83 | NA | 292 | NA |
| **COAD** | 2.92 | NA | 424 | NA |
| **ESCA** | 9.7 | NA | 183 | NA |
| **HNSC** | 20.68 | 6.39 | 523 | 279 |
| **KIRP** | 98.05 | 74.24 | 284 | 161 |
| **LGG** | 113.55 | 247.85 | 510 | 512 |
| **LIHC** | 4.27 | 7.15 | 369 | 193 |
| **LUAD** | 0.64 | 4.51 | 508 | 228 |
| **LUSC** | 3.34 | NA | 488 | NA |
| **MESO** | 0.17 | NA | 86 | NA |
| **OV** | 4.21 | NA | 575 | NA |
| **PAAD** | 15.79 | NA | 182 | NA |
| **READ** | 0.69 | NA | 148 | NA |
| **SARC** | 7.2 | NA | 257 | NA |
| **STAD** | 5.07 | NA | 421 | NA |
| **UCEC** | 28.76 | NA | 526 | NA |
| **UCS** | 0.13 | NA | 56 | NA |
| **UVM** | 22.14 | NA | 80 | NA |

**Supplementary Table 5** – summary of *survClust* **mRNA expression** solution vs available TCGA solutions on logrank (columns 1 and 2). Total number of samples analyzed via *survClust* (column 3), and total samples analyzed via TCGA (column 4)

| Cancer type | survClust logrank | TCGA logrank | survClust(N) | TCGA(N) |
|---|---|---|---|---|
| **BLCA** | 18.51 | 4.8 | 407 | 129 |
| **CESC** | 9.9 | NA | 290 | NA |
| **COAD** | NA | NA | NA | NA |
| **ESCA** | 1.9 | NA | 184 | NA |
| **HNSC** | 25 | 8.73 | 520 | 279 |
| **KIRP** | 93.19 | 20.68 | 288 | 161 |
| **LGG** | 139.33 | 149.71 | 512 | 512 |
| **LIHC** | 26.87 | 4.54 | 366 | 193 |
| **LUAD** | 17.96 | NA | 511 | NA |
| **LUSC** | 5.33 | 0.81 | 500 | 178 |
| **MESO** | 21.24 | NA | 86 | NA |
| **OV** | 5.15 | 4.93 | 303 | 489 |
| **PAAD** | 14.9 | NA | 177 | NA |
| **READ** | 0.87 | NA | 153 | NA |
| **SARC** | 8.62 | NA | 259 | NA |
| **STAD** | 11.66 | NA | 395 | NA |
| **UCEC** | 26.06 | 11.26 | 531 | 537 |
| **UCS** | 2.12 | 0.71 | 56 | 56 |
| **UVM** | 13.95 | NA | 80 | NA |

**Supplementary Table 6** – summary of *survClust* **miRNA expression** solution vs available TCGA solutions on logrank (columns 1 and 2). Total number of samples analyzed via *survClust* (column 3), and total samples analyzed via TCGA (column 4)

| Cancer type | survClust logrank | TCGA logrank | survClust(N) | TCGA(N) |
|---|---|---|---|---|
| **BLCA** | 17.95 | NA | 408 | NA |
| **CESC** | 3.31 | NA | 292 | NA |
| **COAD** | 4.81 | NA | 402 | NA |
| **ESCA** | 4.55 | NA | 182 | NA |
| **HNSC** | 12.89 | 2.52 | 519 | 279 |
| **KIRP** | 26.95 | 8.56 | 284 | 161 |
| **LGG** | 84.91 | NA | 506 | NA |
| **LIHC** | 28.21 | 6.99 | 364 | 193 |
| **LUAD** | 7.28 | NA | 504 | NA |
| **LUSC** | 7.85 | NA | 466 | NA |
| **MESO** | 21.3 | NA | 86 | NA |
| **OV** | 12.47 | NA | 475 | NA |
| **PAAD** | 12.43 | NA | 176 | NA |
| **READ** | 0.13 | NA | 146 | NA |
| **SARC** | 11.36 | NA | 256 | NA |
| **STAD** | 8.76 | NA | 414 | NA |
| **UCEC** | 0.86 | NA | 522 | NA |
| **UCS** | 0.96 | NA | 55 | NA |
| **UVM** | 12.93 | NA | 80 | NA |

**Supplementary Table 7**– summary of *survClust* **Protein expression solution** vs available TCGA solutions on logrank (columns 1 and 2). Total number of samples analyzed via *survClust* (column 3), and total samples analyzed via TCGA (column 4)

| Cancer type | survClust logrank | TCGA logrank | survClust(N) | TCGA(N) |
|---|---|---|---|---|
| **BLCA** | 11.98 | NA | 343 | NA |
| **CESC** | 9.74 | NA | 164 | NA |
| **COAD** | NA | NA | NA | NA |
| **ESCA** | 1.48 | NA | 126 | NA |
| **HNSC** | 17.77 | 5.96 | 212 | 279 |
| **KIRP** | 49.42 | 1.41 | 215 | 161 |
| **LGG** | 53.74 | 14.06 | 425 | 512 |
| **LIHC** | 4.16 | 0.25 | 182 | 193 |
| **LUAD** | 2.26 | NA | 362 | NA |
| **LUSC** | 7.8 | NA | 328 | NA |
| **MESO** | 9.18 | NA | 63 | NA |
| **OV** | 12.62 | NA | 422 | NA |
| **PAAD** | 0.39 | NA | 123 | NA |
| **READ** | NA | NA | NA | NA |
| **SARC** | 18.89 | NA | 223 | NA |
| **STAD** | NA | NA | NA | NA |
| **UCEC** | 17.86 | NA | 439 | NA |
| **UCS** | 0.26 | NA | 47 | NA |
| **UVM** | NA | NA | NA | NA |

**Supplementary Table 8** – Cross tabulation of OV copy number *survClust* labels vs Copy number change in *AKT* gene

| AKT2 | survClust CN labels c1 | c2 | c3 | c4 | c5 | c6 |
|---|---|---|---|---|---|---|
| **-2** | 0 | 0 | 1 | 1 | 0 | 0 |
| **-1** | 26 | 14 | 84 | 30 | 3 | 31 |
| **0** | 76 | 46 | 3 | 43 | 1 | 4 |
| **1** | 49 | 42 | 1 | 29 | 31 | 10 |
| **2** | 4 | 7 | 0 | 3 | 18 | 1 |

### Supplementary figure 1

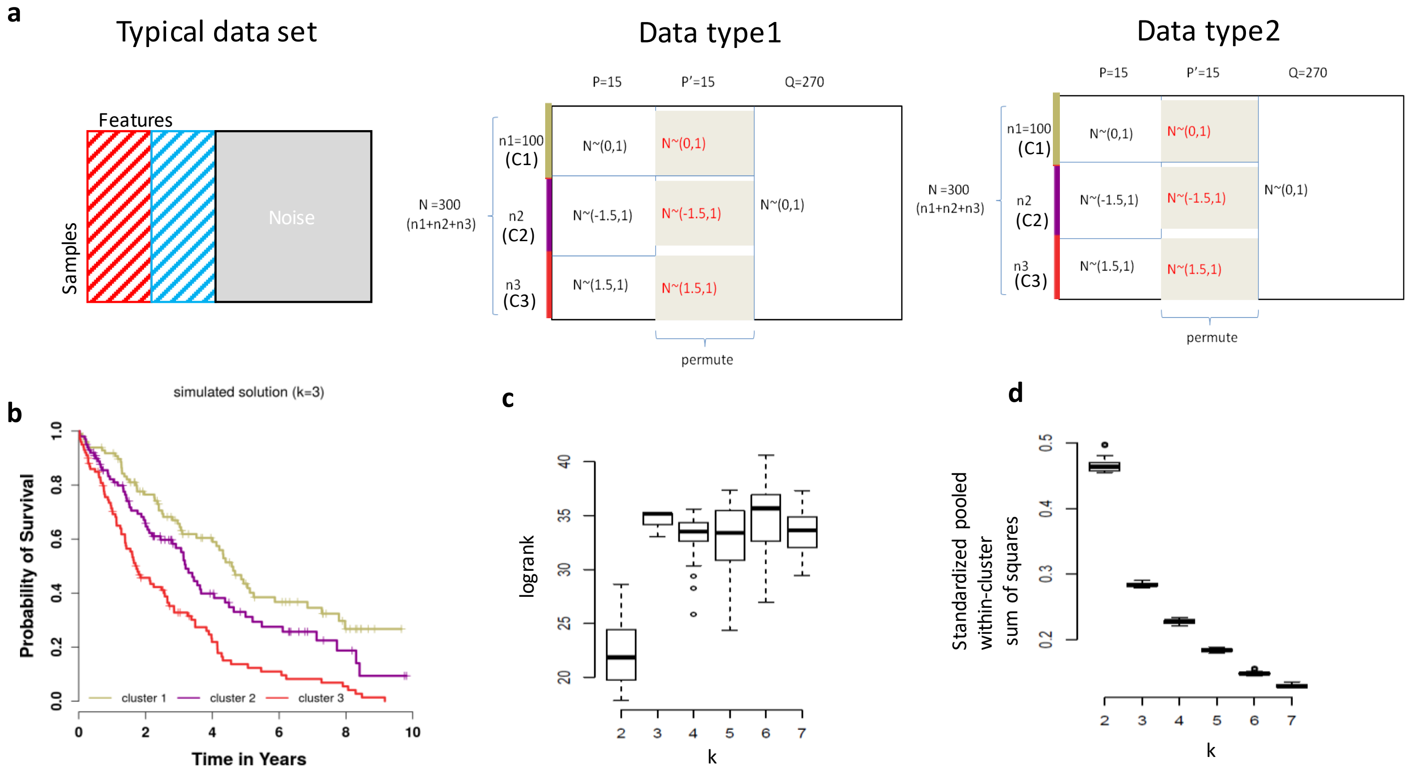


e

| survClust labels | c1 | c2 | c3 |
|---|---|---|---|
| Simulated labels | | | |
| Cluster 1 | 100 | 0 | 0 |
| Cluster 2 | 0 | 100 | 0 |
| Cluster 3 | 0 | 0 | 100 |

**Supplementary figure 1. Simulation analysis.**
**(a)** A typical data set with survival related features, survival unrelated features and noise. **Data Type 1** – 15 out of 300 features have a distinct structure along with survival association, next 15 were simulated in the same manner but permutated, and remaining features were noise. **Data Type 2 –** Same as Data type 1.

**(b)** Simulated 3-class survival with median survival time as 4,3, and 2 years in clusters 1,2 and 3 respectively.

**(c)** Boxplot of logrank of 50 rounds of cross validation with *survClust.*

**(d)** Boxplot of standardized pooled within-cluster sum of squares of 50 rounds of cross validation with *survClust.*

**(e)** Cross tabulation with *survClust* labels with respect to simulated class labels or truth.

### Supplementary figure 2

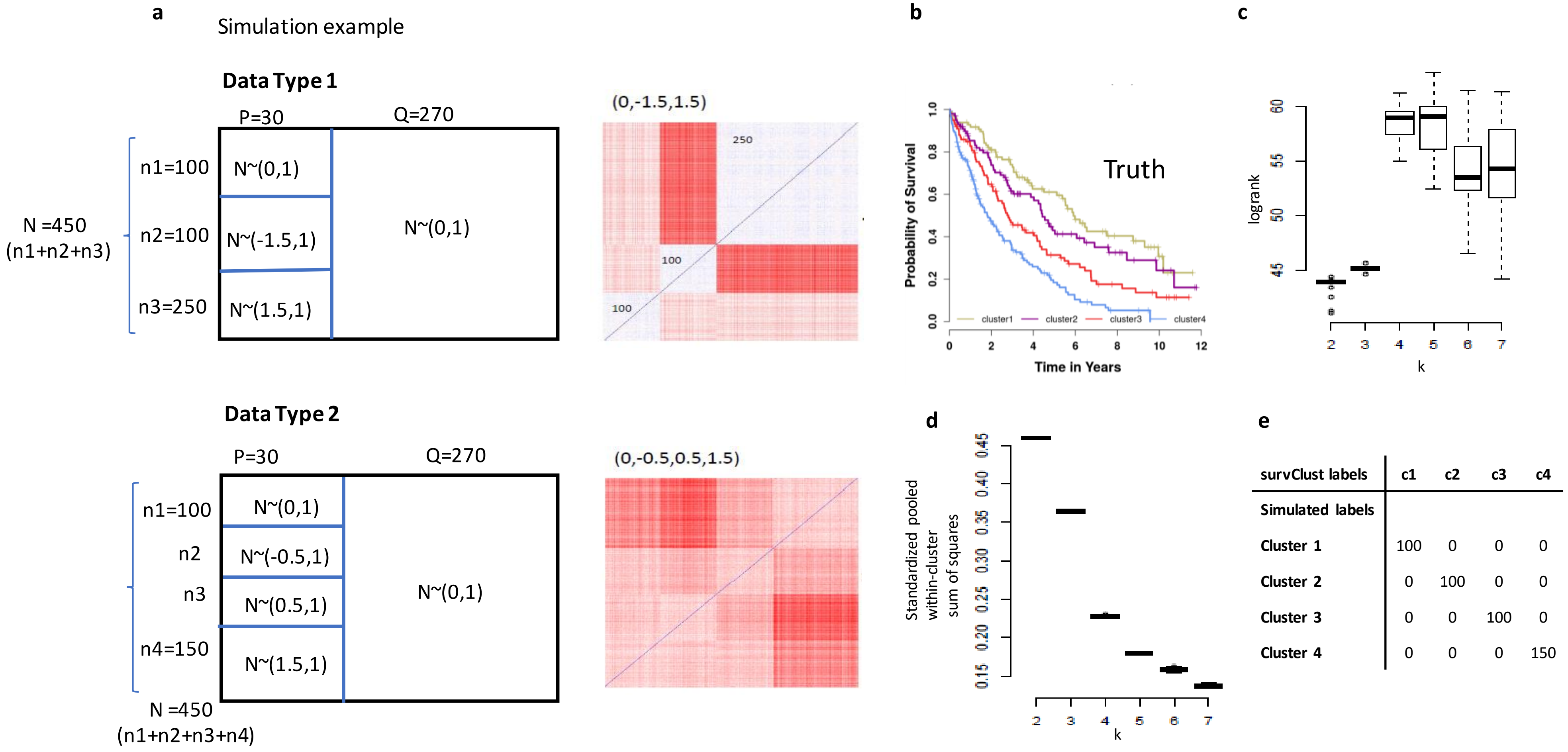


| survClust labels | c1 | c2 | c3 | c4 |
|---|---|---|---|---|
| Simulated labels | | | | |
| Cluster 1 | 100 | 0 | 0 | 0 |
| Cluster 2 | 0 | 100 | 0 | 0 |
| Cluster 3 | 0 | 0 | 100 | 0 |
| Cluster 4 | 0 | 0 | 0 | 150 |

**Supplementary figure 2. Second simulation analysis.**
**(a)** Simulated structures of two data types. **Data type 1** – strong clusters, weak survival association. Plot on the right is the weighted distance matrix of data type 1. **Data type 2** – weak clusters, strong cluster association. Plot on the right is the weighted distance matrix of data type 2.

**(b)** Simulated 4-class survival with median survival time as 5,4,3, and 2 years in clusters 1,2,3 and 4 respectively.

**(c)** Boxplot of logrank of 50 rounds of cross validation with *survClust.*

**(d)** Boxplot of standardized pooled within-cluster sum of squares of 50 rounds of cross validation with *survClust.*

**(e)** Cross tabulation with *survClust* labels with respect to simulated class labels or truth.

**Supplementary Figure 3**

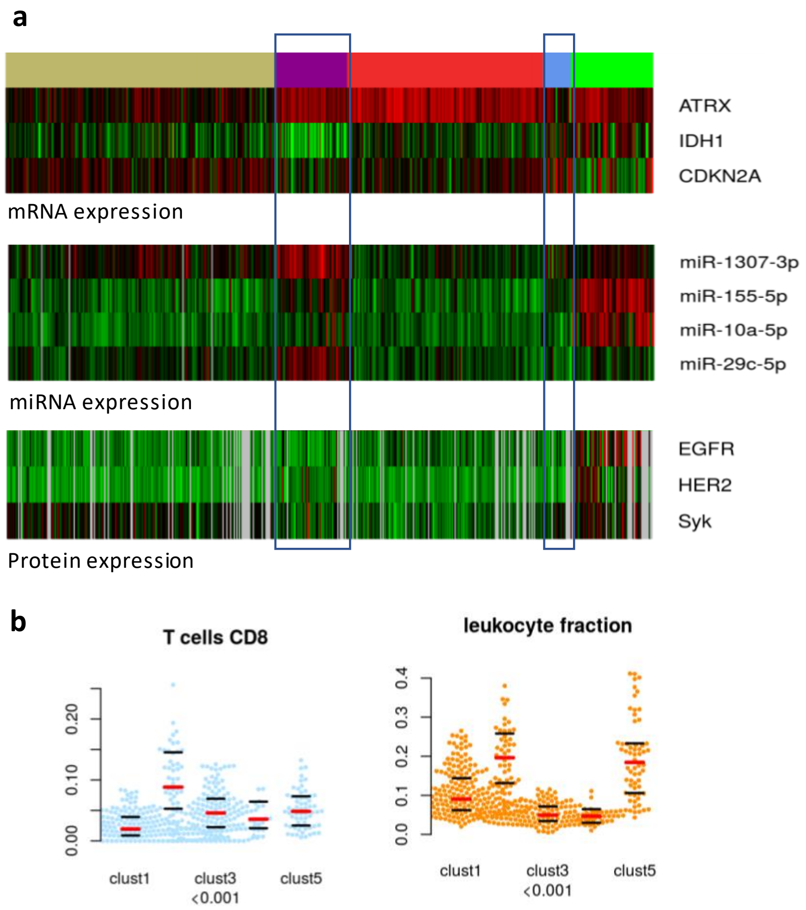


**Supplementary Figure 3: *survClust* integrated solution of LGG**

**(a)** Selected gene, miRNA and protein expression with respect to 5-class LGG integrated *survClust* labels. **(b)** Beeswarm plots summarizing infiltration levels (y-axis) of CD8 T-cells and leukocyte fraction across the five groups (x-axis). Red line depicts the median, and top and bottom black bars represent 25th and 75th percentile respectively.

### Supplementary Figure 4

Mutation - Yes
Missing
HPV negative
HPV Positive
Yes
No

c1 c3 c5
c2 c4 c6

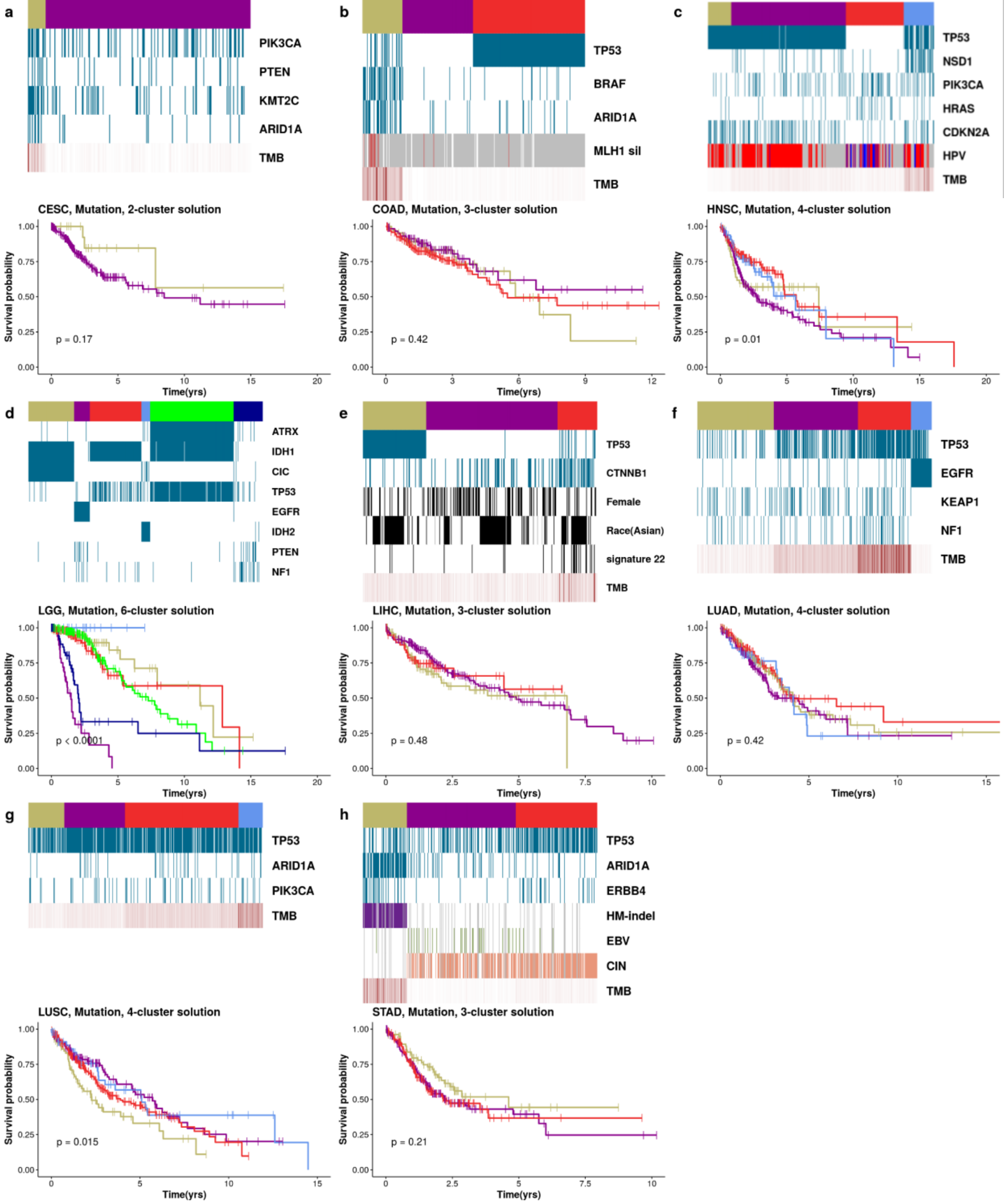


**Supplementary Figure 4: *survClust* identifies TMB patterns across cancer types**

Each sub figure contains *panelmap* summarizing differentiating molecular and clinical characteristics for *k* cluster *survClust* solution labels on mutation data alone, followed by Kaplan-Meier curves for each group. **(a)**CESC **(b)**COAD **(c)**HNSC **(d)**LGG **(e)**LIHC **(f)**LUAD **(g)**LUSC **(h)**STAD

### Supplementary Figure 5

**Supplementary Figure 5: CD8 T-cell distribution of *survClust* mutation classes of various cancer types**

Boxplot summarizing CD8 T-cell expression (y-axis) across survClust class labels (x-axis). Red line depicts the median, and top and bottom black bars represent 25th and 75th percentile respectively. Significance from association test is shown at the bottom of each plot.
**(a)**CESC **(b)**COAD **(c)**HNSC **(d)**LGG **(e)**LIHC **(f)**LUAD **(g)**LUSC **(h)**STAD

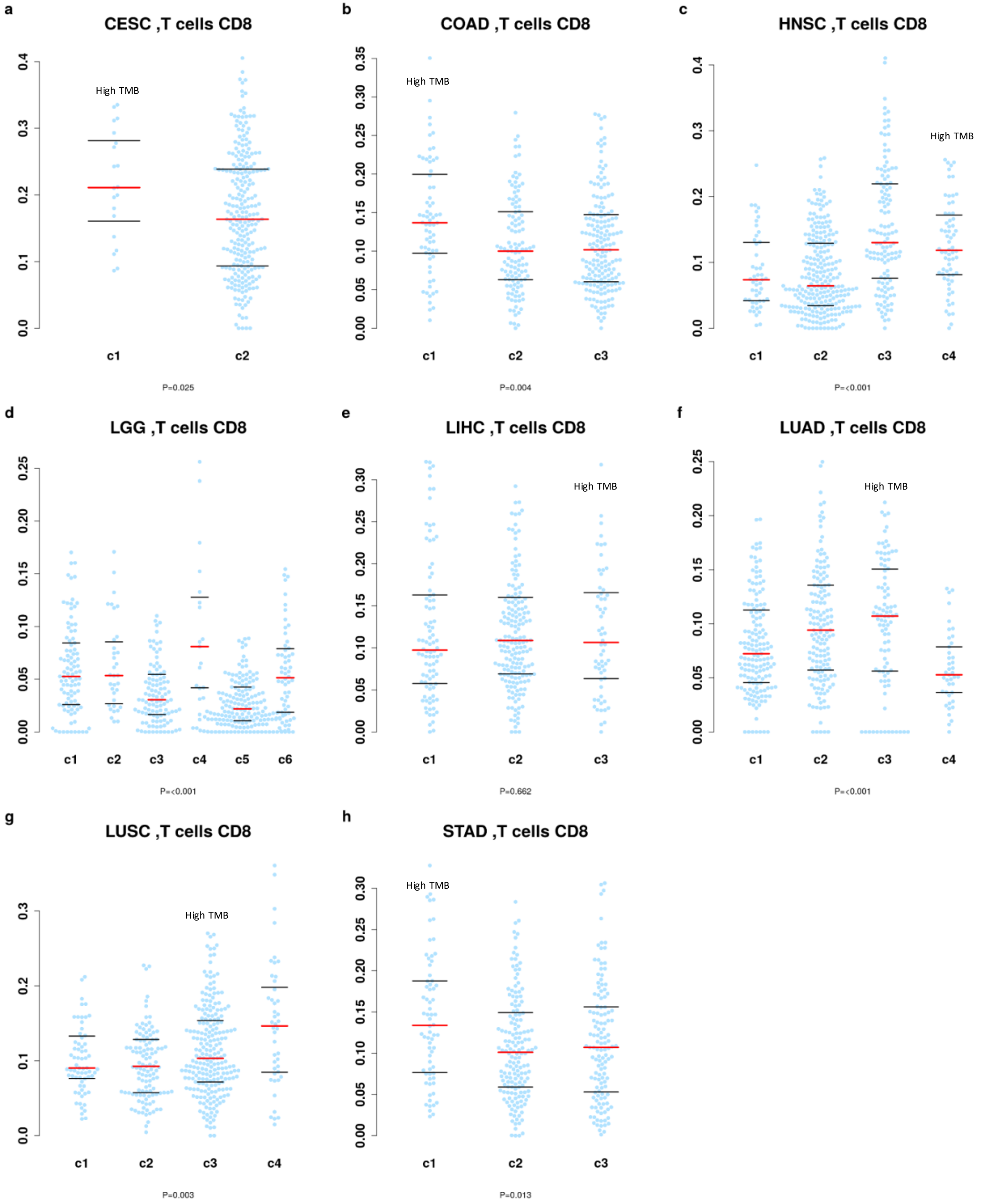

### Supplementary Figure 6

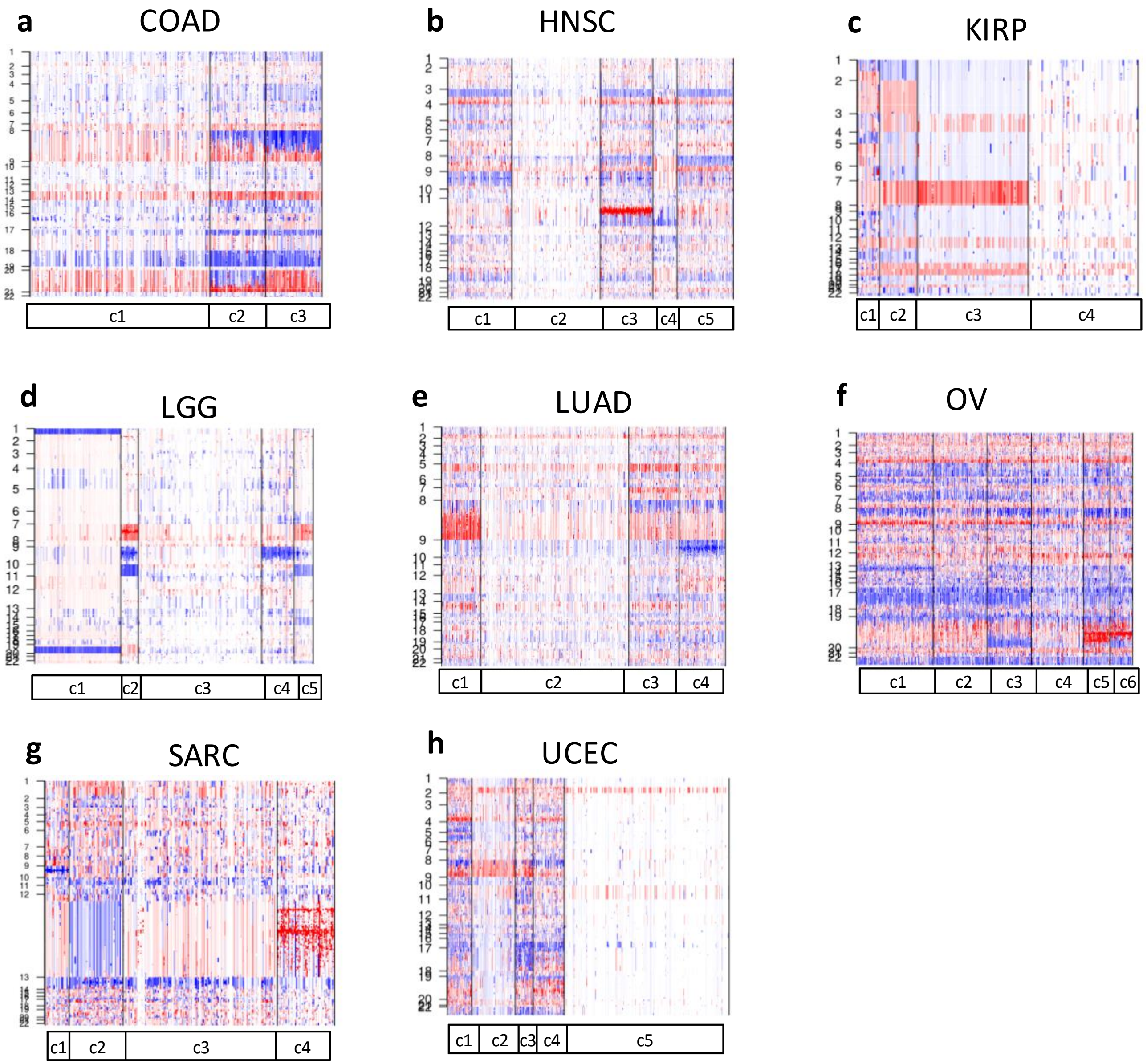


**Supplementary Figure 6: *survClust* identifies FGA patterns across cancer types: Global view of Copy Number Change**

Figures a-i represent a global view of copy number changes (Chromosome 1-22) in cluster labels identified by survClust run on segmented copy number data. **(a)**COAD **(b)**HNSC **(c)**KIRP **(d)**LGG **(e)**LUAD **(f)**OV **(g)**SARC **(h)**UCEC.

### Supplementary Figure 7

**Supplementary Figure 7: *survClust* identifies FGA patterns across cancer types: Kaplan-Meier Curves**

Kaplan-Meier curves for each group obtained by *survClust* run on segmented copy number data for various cancer types.
**(a)**COAD **(b)**HNSC **(c)**KIRP **(d)**LGG **(e)**LIHC **(f)**LUAD **(g)**OV **(h)**SARC **(i)**UCEC

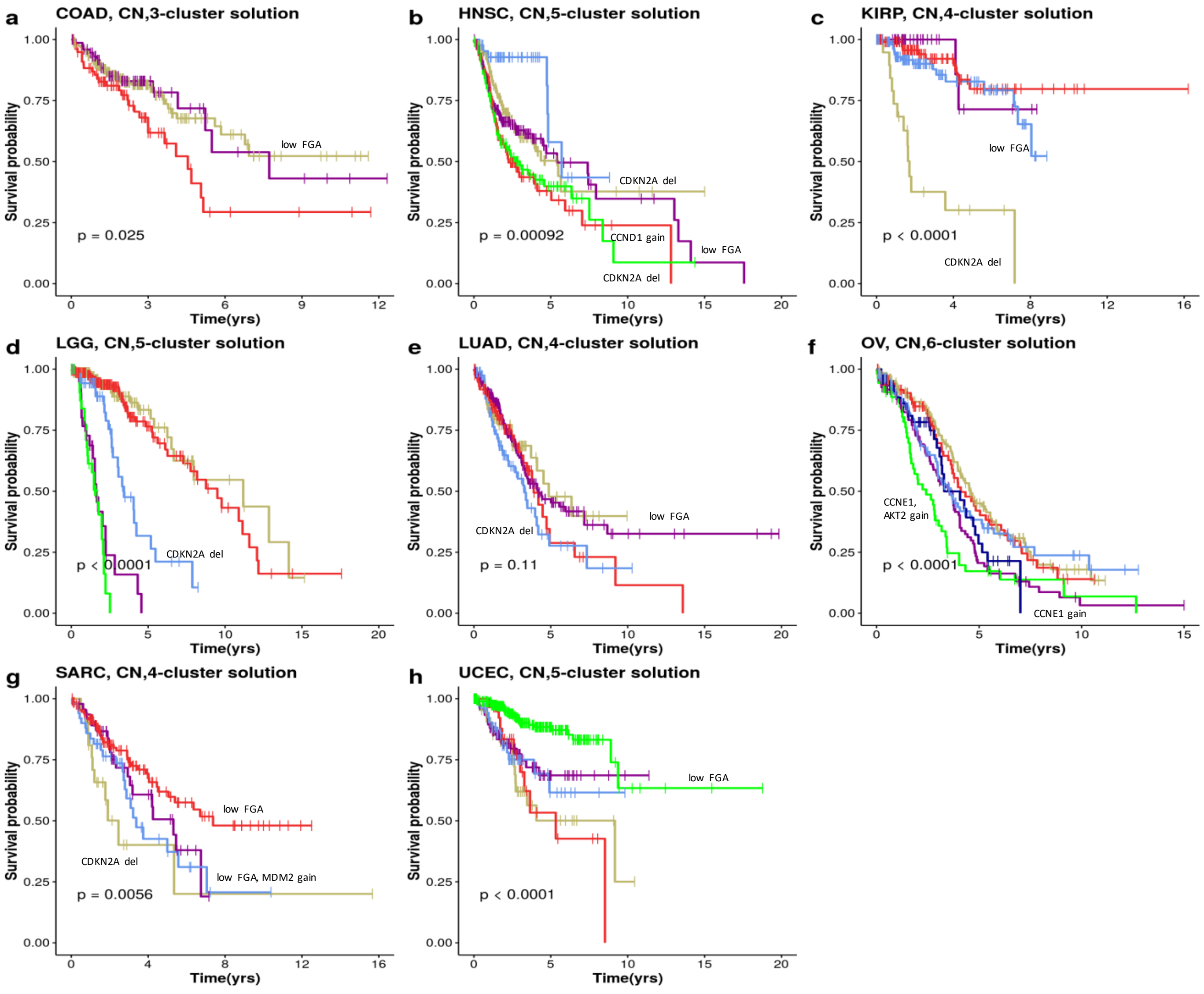


CCNE! gain

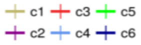

### Supplementary Figure 8

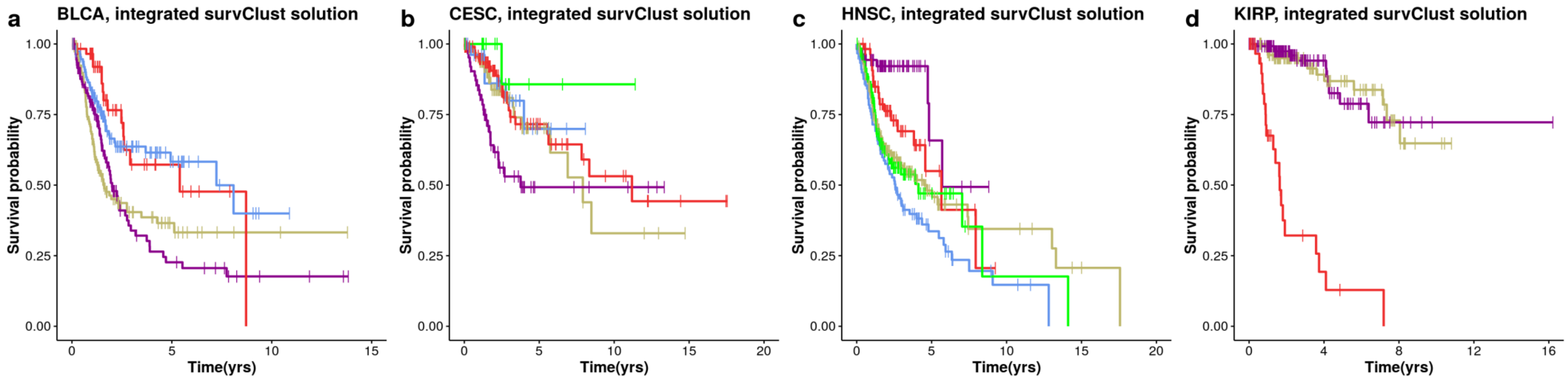


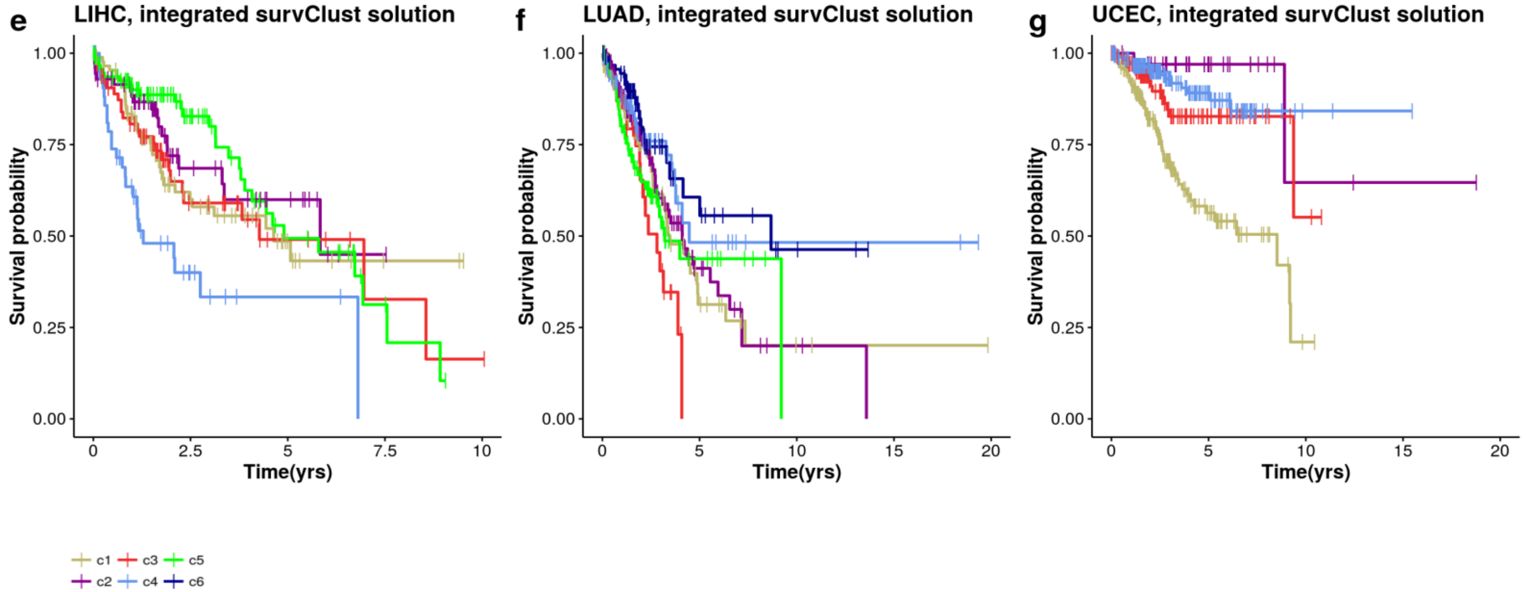


**Supplementary Figure 8: *survClust* integrated solution of various cancer types**

Kaplan-Meier curves for solutions obtained by *survClust* run on integrating six platforms – DNA methylation, Copy Number, mRNA expression, miRNA expression, protein expression and mutation data.
**(a)**BLCA **(b)**CESC**(c)**HNSC **(d)**LIHC**(e)**KIRP**(f)**LUAD **(g)**CESC
